# A reference map of the human protein interactome

**DOI:** 10.1101/605451

**Authors:** Katja Luck, Dae-Kyum Kim, Luke Lambourne, Kerstin Spirohn, Bridget E. Begg, Wenting Bian, Ruth Brignall, Tiziana Cafarelli, Francisco J. Campos-Laborie, Benoit Charloteaux, Dongsic Choi, Atina G. Cote, Meaghan Daley, Steven Deimling, Alice Desbuleux, Amélie Dricot, Marinella Gebbia, Madeleine F. Hardy, Nishka Kishore, Jennifer J. Knapp, István A. Kovács, Irma Lemmens, Miles W. Mee, Joseph C. Mellor, Carl Pollis, Carles Pons, Aaron D. Richardson, Sadie Schlabach, Bridget Teeking, Anupama Yadav, Mariana Babor, Dawit Balcha, Omer Basha, Christian Bowman-Colin, Suet-Feung Chin, Soon Gang Choi, Claudia Colabella, Georges Coppin, Cassandra D’Amata, David De Ridder, Steffi De Rouck, Miquel Duran-Frigola, Hanane Ennajdaoui, Florian Goebels, Liana Goehring, Anjali Gopal, Ghazal Haddad, Elodie Hatchi, Mohamed Helmy, Yves Jacob, Yoseph Kassa, Serena Landini, Roujia Li, Natascha van Lieshout, Andrew MacWilliams, Dylan Markey, Joseph N. Paulson, Sudharshan Rangarajan, John Rasla, Ashyad Rayhan, Thomas Rolland, Adriana San-Miguel, Yun Shen, Dayag Sheykhkarimli, Gloria M. Sheynkman, Eyal Simonovsky, Murat Taşan, Alexander Tejeda, Jean-Claude Twizere, Yang Wang, Robert J. Weatheritt, Jochen Weile, Yu Xia, Xinping Yang, Esti Yeger-Lotem, Quan Zhong, Patrick Aloy, Gary D. Bader, Javier De Las Rivas, Suzanne Gaudet, Tong Hao, Janusz Rak, Jan Tavernier, Vincent Tropepe, David E. Hill, Marc Vidal, Frederick P. Roth, Michael A. Calderwood

## Abstract

Global insights into cellular organization and function require comprehensive understanding of interactome networks. Similar to how a reference genome sequence revolutionized human genetics, a reference map of the human interactome network is critical to fully understand genotype-phenotype relationships. Here we present the first human “all-by-all” binary reference interactome map, or “HuRI”. With ~53,000 high-quality protein-protein interactions (PPIs), HuRI is approximately four times larger than the information curated from small-scale studies available in the literature. Integrating HuRI with genome, transcriptome and proteome data enables the study of cellular function within essentially any physiological or pathological cellular context. We demonstrate the use of HuRI in identifying specific subcellular roles of PPIs and protein function modulation via splicing during brain development. Inferred tissue-specific networks reveal general principles for the formation of cellular context-specific functions and elucidate potential molecular mechanisms underlying tissue-specific phenotypes of Mendelian diseases. HuRI thus represents an unprecedented, systematic reference linking genomic variation to phenotypic outcomes.

---

The reference human genome sequence has enabled systematic study of genetic^1^ and expression^2^ variability at the organism^1^, tissue^2^, cell type^3^ and single cell level^4^. Despite advances in sequencing genomes, transcriptomes and proteomes, we still understand little of the cellular mechanisms that mediate phenotypic and tissue or cell type variability. A mechanistic understanding of cellular function and organization emerges from studying how genes and their products, primarily proteins, interact with each other, forming a dynamic interactome that drives biological function. Analogous to the reference human genome sequence^5,6^, a reference map of the human protein interactome, generated systematically and comprehensively, would provide an unprecedented scaffold for the unbiased proteome-wide study of biological mechanisms, generally and within specific cellular contexts. Almost 20 years after the publication of a first draft of the reference human genome sequence^5,6^, a reference protein interactome map is yet to be reported.

Proteins are biochemically more complex than DNA, the interactome is much more dynamic than the genome, and the search space for interactions requires testing all-by-all pairwise combinations, making interactome mapping extremely challenging. Approaches to human proteome-wide protein-protein interaction (PPI) mapping either aim to identify protein complex assemblies using mass spectrometry^7–9^ or direct PPIs using binary screening methods such as yeast two-hybrid (Y2H) followed by empirical validation using orthogonal assays^10–12^. In contrast to protein complex mapping, binary interactome mapping is based on interrogating pairs of proteins for interaction independently from any particular endogenous cellular context, thereby generating relatively unbiased systematic PPI datasets. For example, our most recent human protein interactome map (HI-II-14) described ~14,000 PPIs involving 4,000 proteins from screening ~40% of the genome-by-genome search space^10^ and in striking contrast to literature-curated and protein complex interactome maps, HI-II-14 uniformly covered the proteome, free of study and expression bias.

To increase interactome coverage and generate a first human reference interactome map, we have expanded the ORFeome collection to encompass ~90% of the protein-coding genome and screened this search space a total of nine times with a panel of assays using an enhanced screening platform. The resulting human binary PPI map doubles HI-II-14’s coverage of the proteome and quadruples its interactome coverage. Integrating this PPI network with genome, transcriptome and proteome resources, we infer cellular context-specific views of the protein interactome, which are predictive of cellular context-specific gene function, at the level of individual subcellular compartments, cell types and tissues, across developmental stages and in disease (Fig. 1a). With its comprehensive view of the protein interactome, the resulting network enables biological discovery across any cellular context, thus representing the first reference map of the human protein interactome.

**Fig. 1.**
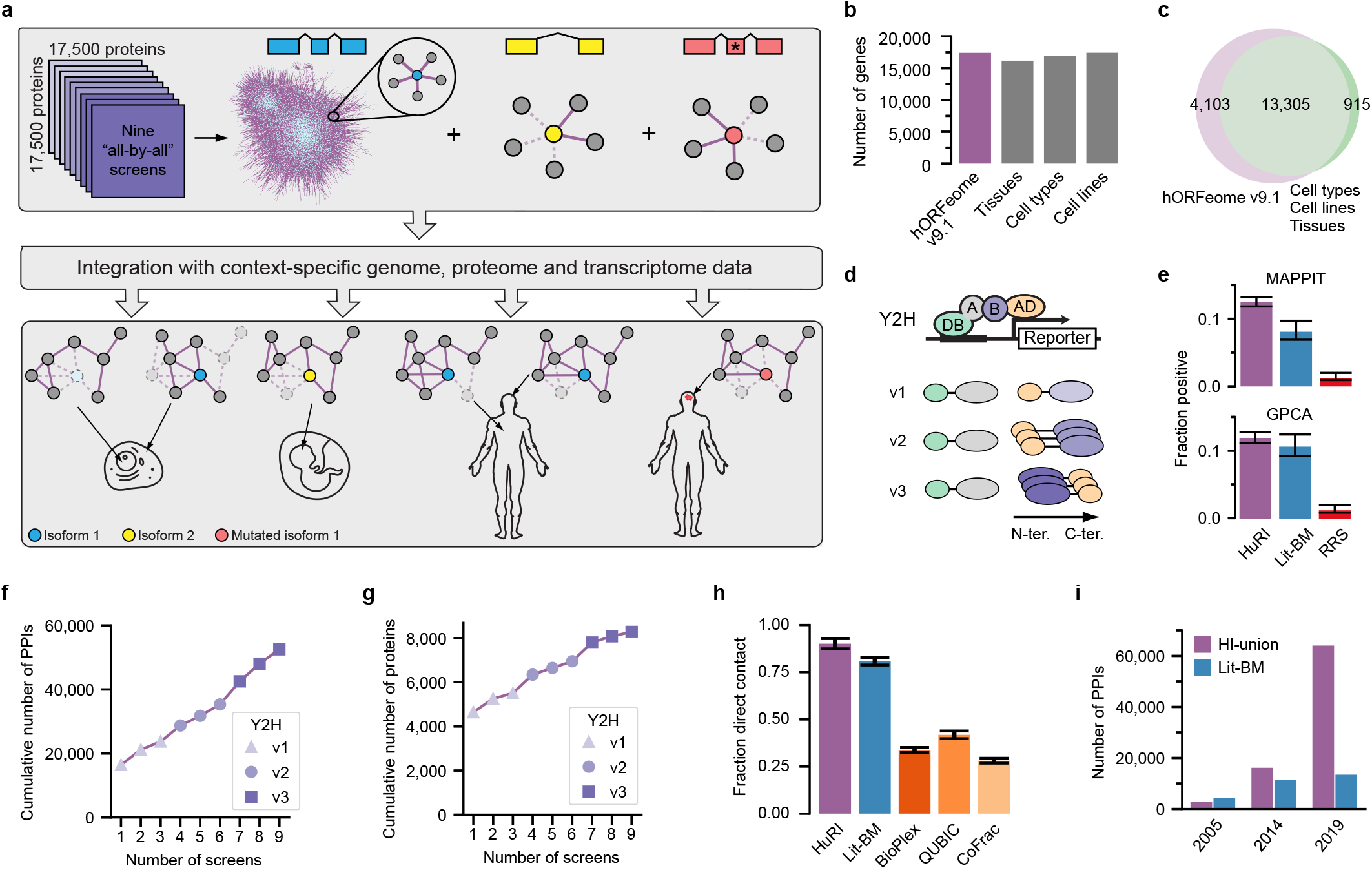
Generation of a reference interactome map using a panel of binary assays. **a**, Overview of HuRI generation and applications. **b**, Number of protein-coding genes in hORFeome v9.1 and GTEx, FANTOM, and HPA transcriptome projects. **c**, Overlap between hORFeome v9.1 and intersection of transcriptomes in **b. d**, Schematic of the Y2H assay versions. **e**, Experimental validation. PPIs were tested in GPCA (first six screens) and MAPPIT (all nine screens). Lit-BM: literature-curated binary PPIs with multiple evidence; RRS: random protein pairs. Error bars are 68.3% Bayesian CI. **f, g**, Number of PPIs and proteins, detected with each additional screen. **h**, Fraction of direct contact pairs among five PPI networks. Error bars are standard error of proportion. **i**, Number of PPIs identified over time from screening at CCSB and Lit-BM.

## Generation and biophysical characterization of HI-III-19

Our previously published human protein interactome map, HI-II-14^10^, covered less than half of the possible search space. To generate a more complete map, we established human ORFeome v9.1. This expanded ORFeome covers 17,408 protein-coding genes, on par with the number of genes found to be expressed in three comprehensive individual transcriptome sequencing studies^2,3,13^ (Fig. 1b) and includes 94% of the genes with robust evidence of expression in all three (Fig. 1c, Supplementary Table 1). The search space formed by hORFeome v9.1 (Space III), encompassing over 150 million binary combinations, more than doubles the space screened to generate HI-II-14 and represents the most comprehensive search space to be systematically screened for human PPIs.

Limitations in PPI assay sensitivity can be overcome by employing different PPI assays^14^ or different versions of the same PPI assay^15,16^. To maximize sensitivity while maintaining high-throughput screening capabilities, we employed three Y2H assay versions (Fig. 1d), which, when benchmarked against a gold standard positive and random reference set (PRSv1 and RRSv1)^17^, showed good sensitivity and low false positive rates while detecting complementary sets of PPIs (Extended Data Fig. 1a, b, Supplementary Table 2). We further assessed Y2H assay version quality, complementarity and screening behavior on a test space of ~2,000 by ~2,000 human genes^10^. After verification by pairwise Y2H retesting and sequence confirmation, PPIs from each version were evaluated using MAPPIT^18^, an orthogonal assay. For each Y2H assay version, the recovery rate of PPIs was comparable or exceeded that of a set of PPIs from the literature with ≥ 2 pieces of experimental evidence, of which at least one comes from a binary assay type (Lit-BM)^10^ (Extended Data Fig. 1c, Supplementary Table 3). The Y2H assay versions were complementary, in that performing three screens with each version doubled the number of PPIs and proteins detected relative to performing the equivalent number of screens using a single assay version (Extended Data Fig. 1d, Supplementary Table 4).

To construct the reference interactome, we performed nine screens of Space III, three with each Y2H assay version, followed by pairwise testing in quadruplicate, sequence confirmation, and validation using two orthogonal assays, MAPPIT^18^ and GPCA^19^. Screen PPIs were recovered at rates that were similar or superior to Lit-BM over a large range of score thresholds (Fig. 1e, Extended Data Fig. 1e-g, Supplementary Table 5), confirming the high quality of the interactome dataset. Each additional screen identified novel PPIs and proteins, with the largest gains obtained by switching assay versions (Fig. 1f, g, Extended Data Fig. 1d), highlighting the importance of performing multiple screens and using several assay versions. The dataset, versioned HI-III-19 (Human Interactome obtained from screening Space III, published in 2019), contains 52,569 verified PPIs involving 8,275 proteins (Supplementary Table 6). Given its systematic nature, completeness and scale, we consider HI-III-19 to be the first draft of the Human Reference Interactome (HuRI).

To assess whether assay complementarity can partially stem from different steric constraints in the protein fusions, we integrated HuRI with protein structure data^20^ and observed that the dataset is depleted for PPIs where the interaction interface is a short spatial distance (<20Å) from the protein terminus fused to the AD domain (Extended Data Fig. 2a, b, Supplementary Table 7). These results provide the first systematic investigation into the impact of protein tags on PPI detection.

Molecular mechanisms can be more readily inferred from direct PPIs, yet, the fraction of direct versus indirect PPIs reported in various human protein interactome maps is unknown. Using three-dimensional structural information from protein complexes with at least three subunits^20,21^, we show that the vast majority of PPIs in HuRI (90%) correspond to direct biophysical contacts, significantly higher than in Lit-BM (81%, *P* = 0.019, two-sided Fisher’s exact test, *n* = 121 (HuRI), 410 (Lit-BM)) or in protein complex interactome maps (less than 50%, *P* < 0.001 for all, two-sided Fisher’s exact test) (Fig. 1h, Supplementary Table 8), demonstrating that HuRI represents a unique dataset of direct PPIs. Combining HuRI with all previously published systematic screening efforts at CCSB yields 64,006 binary PPIs involving 9,094 proteins (HI-union) (Supplementary Table 9), which is approximately five-fold more PPIs than the entirety of high-quality binary PPIs curated from the literature (Fig. 1i). The union of Lit-BM and HI-union represent the most complete collection of high quality direct PPI data available to date (http://interactome.dfci.harvard.edu/huri/).

PPIs in HuRI vary by the number of screens and assay versions in which they were detected (Extended Data Fig. 2c, d). To investigate any potential relationship between these factors and PPI false discovery rate, we compared MAPPIT recovery rates of HuRI and Lit-BM PPIs found in different numbers of screens. Interestingly, both sets of PPIs show that MAPPIT recovery rates increase with the number of screens in which an interaction was detected (Extended Data Fig. 2e, Supplementary Table 10). This trend persists even when titrating Lit-BM to higher numbers of experimental evidence (Extended Data Fig. 2f), suggesting that differences in PPI recovery rates are driven by factors other than veracity of a PPI. In HuRI the number of screens in which an interaction is detected is weakly correlated with both the size of the molecular interfacial area (*ρ* = 0.15, *P* = 0.026, two-sided permutation test, *n* = 234) (Extended Data Fig. 2g) and the number of atomic contacts (*ρ* = 0.14, *P* = 0.038, two-sided permutation test, *n* = 234) (Extended Data Fig. 2h, Supplementary Table 11), suggesting that identification of a PPI in a screen is impacted by interaction strength and may therefore be reflecting ‘detectability’ rather than accuracy. Indeed, PPIs in HuRI found in at least two screens corresponded more often to direct PPIs within rather than between well-described stable protein complexes^22,23^ (*P* = 3 × 10^-18^, two-sided Fisher’s exact test, *n* = 1817) (Extended Data Fig. 2i, 3, Supplementary Table 12). Because the majority of PPIs in HuRI were found in only one screen, our data further reinforces previous observations^7,24^ that the protein interactome might be dominated by weak, more transient PPIs, that are harder to detect. PPI detectability may impact previous assessments of overlap between PPI datasets, as well as estimates of interactome size.

## Multiple layers of functional relationships between proteins in HuRI

Based on the observation that HuRI is enriched in direct PPIs, we hypothesize that proteins in HuRI with similar interaction interfaces should share a significant number of their interaction partners. For example, retinoic acid receptors RXR-γ and -β (Fig. 2a, left panel) share previously reported interaction partners involving binding to retinoic acid receptor RAR types^25^ and oxysterol receptors^26^. We derived a profile similarity network (PSN) from HuRI (Supplementary Table 13), and found that the number of pairs of proteins in HuRI with similar interaction profiles is significantly higher than random (*P* < 0.01, one-sided empirical test) (Extended Data Fig. 4a) and proteins of overall higher sequence identity tend to exhibit higher interaction profile similarities (*P* < 0.01, one-sided empirical test) (Extended Data Fig. 4b). However, proteins with a tendency to share interaction partners often have interaction interfaces that are similar, as opposed to complementary, and therefore tend not to interact, unless both proteins originate from the same ancestral protein that was able to self-interact^27^ (Extended Data Fig. 4c). Indeed, only 5% of the proteins found to interact in HuRI share more than 10% of their interaction partners. HuRI and the PSN display significant enrichment to link proteins of similar function (*P* < 0.01, one-sided empirical test) (Fig. 2b, Extended Data Fig. 4d, e) and both contain much higher numbers of functional modules^28^ compared to our previously published interactome maps^10,11^ (Fig. 2c). Because HuRI and the PSN display little link overlap but both are functionally enriched, this suggests that HuRI and the PSN complement each other in revealing functional relationships between proteins.

**Fig. 2.**
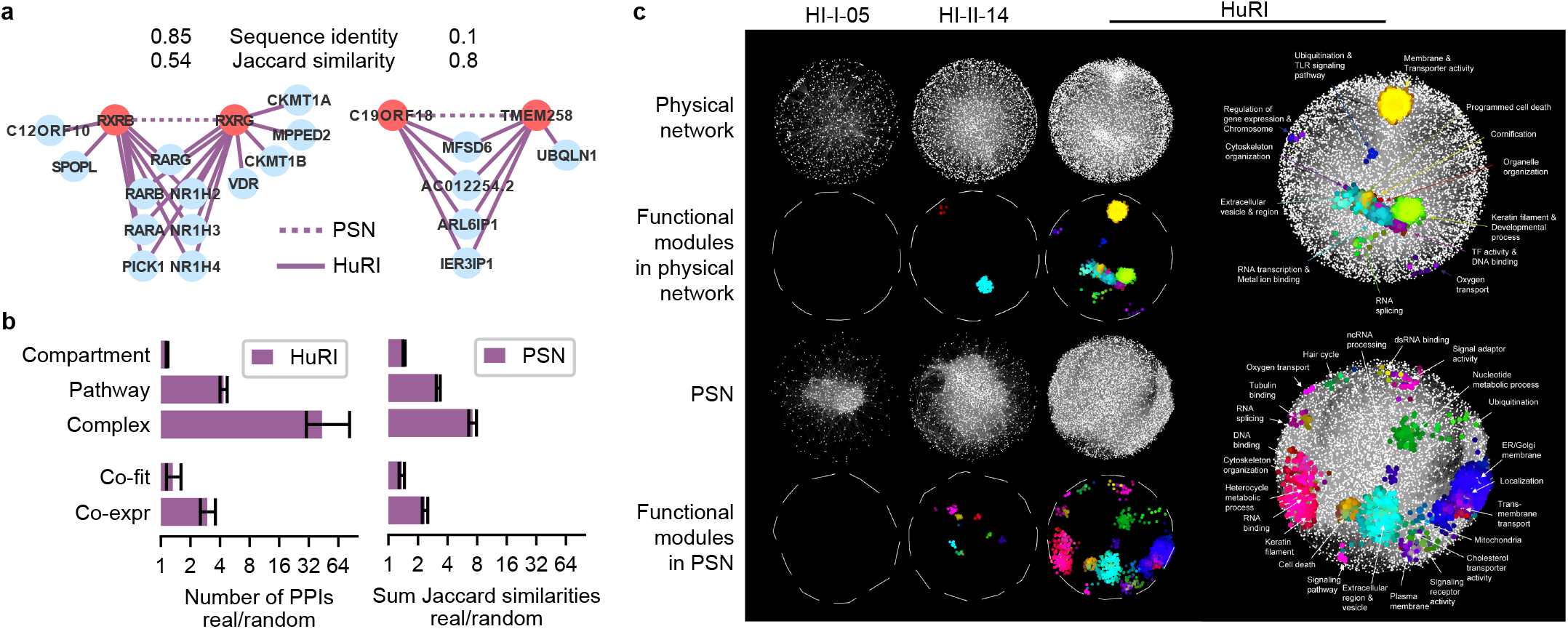
Complementary functional relationships in HuRI between genes originating from interaction profile similarity and PPIs. **a**, Examples of protein pairs in HuRI with high interaction profile similarity and both high (left) and low sequence identity (right). **b**, Fold-enrichment of HuRI and its profile similarity network (PSN) for protein pairs with shared functional annotation. Error bars are 95% confidence intervals. **c**, Functional modules in HuRI and its PSN and in previously published interactome maps from CCSB.

As shown above, global sequence identity between two proteins is indicative of shared interaction interfaces, however, it likely fails to identify pairs of proteins whose shared interaction interface is small. Indeed, 50% (502) of all pairs of proteins in HuRI with interaction profile similarities ≥ 0.5 exhibit less than 20% sequence identity, showing that the functional relationships between proteins cannot necessarily be identified by sequence identity. One such pair of proteins is the endoplasmic reticulum (ER) transmembrane protein TMEM258 and the uncharacterized protein C19ORF18, which display a sequence identity of only 10% but share 80% of their interactors (Fig. 2a, right panel). TMEM258 catalyzes the first step in N-glycosylation of proteins in the ER and might play a role in protein translocation across the ER^29^. Roles in protein transport and ER function have also been ascribed to two of the four shared interaction partners, ARL6IP1^30^ and IER3IP1^31^, suggesting that C19ORF18 as well as the other two shared yet unstudied interaction partners MFSD6 and AC012254.2 might contribute to ER-related functions of protein maturation and transport.

## Uncharted network neighborhoods of disease-related genes

Unlike Lit-BM, HuRI was generated by systematically testing pairs of proteins for interaction. While Lit-BM is highly biased towards the most highly studied genes^10^, HuRI covers the genome-by-genome space, as ranked by number of publications, more uniformly and at increased depth compared to Lit-BM and our previous screening efforts (Fig. 3a). Considering these differences in interactome space coverage, we find that among the best-studied genes, where Lit-BM is most complete, the agreement between Lit-BM and HuRI is highest, with ~40% of the PPIs in HuRI being previously identified (Fig. 3b). Because of its uniform coverage, HuRI substantially expands the set of genes of biomedical interest for which high-quality direct PPI data is available (Fig. 3c, Extended Data Fig. 4g), and extends their network neighborhood to previously uncharted regions of the protein interactome (Fig. 3d, Extended Data Fig. 4g).

**Fig. 3.**
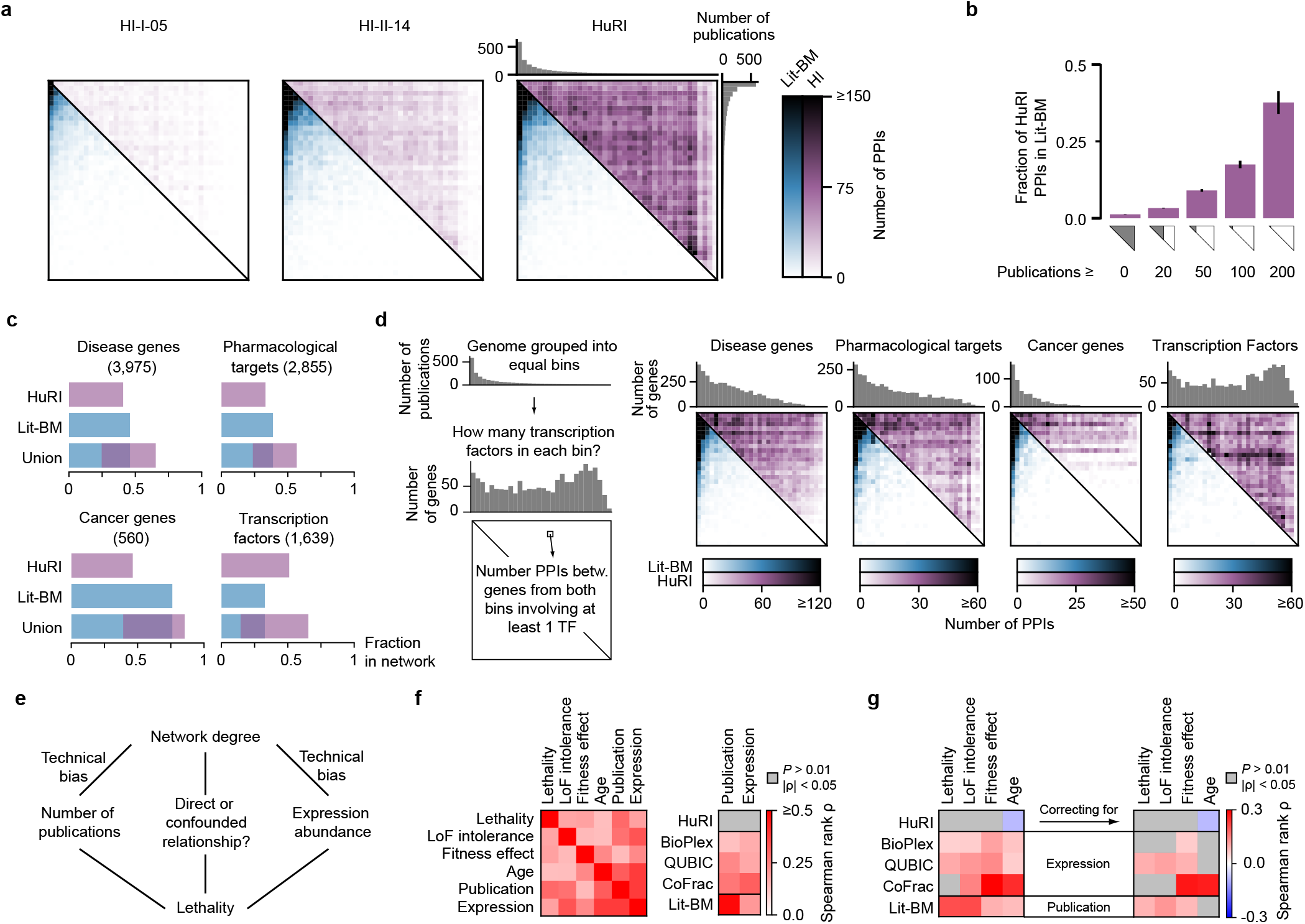
Unbiased proteome coverage of HuRI reveals uncharted network neighborhoods of disease-related genes. **a**, Heatmaps of Y2H PPI counts, ordered by number of publications, for CCSB datasets and Lit-BM. **b**, Fraction of HuRI PPIs in Lit-BM, for increasing values of the minimum number of publications per protein. Error bars are standard error of proportion. **c**, Fraction of genes with at least one PPI for biomedically interesting genes. **d**, As **a**, but restricted to PPIs involving genes from the corresponding gene sets. **e**, Schematic of relation between variables: observed PPI degree, abundance, study bias and lethality. **f**, Correlation matrices. LoF: Loss-of-Function. PPI datasets refer to their network degree. **g**, Correlation between degree and variables of interest, before (left) and after (right) correcting for the technical confounding factors.

As previously shown^10,32^, study bias can skew interactome coverage and the assessment of systems properties of genes. Using HuRI, we find no evidence of reported correlations between a protein’s number of interaction partners (degree) and various gene properties, i.e. lethality^33,34^, loss-of-function intolerance^1^, fitness effect^35^, and age^36,37^ (Fig. 3e-g). Moreover, these correlations weaken for protein complex and Lit-BM interactome maps when they are corrected for confounding protein expression or study bias, respectively (Fig. 3e-g). These results highlight the value of HuRI as a uniformly-mapped reference for the study of systems properties of genes and networks.

## Identification of subcellular compartment-specific roles of PPIs

Proteins are localized to specific subcellular compartments to exert functions that can depend both on the subcellular environment and the local PPI network. Despite available proteome-wide datasets on the localization of individual proteins^38^, experimental determination of cellular localization-specific PPI networks remains challenging^39^. We find that proteins localized to a diverse range of subcellular compartments are evenly represented in HuRI (Fig. 4a) suggesting that cellular localization-specific PPI networks can be inferred for many different cellular compartments via integration of HuRI with available protein localization data.

**Fig. 4.**
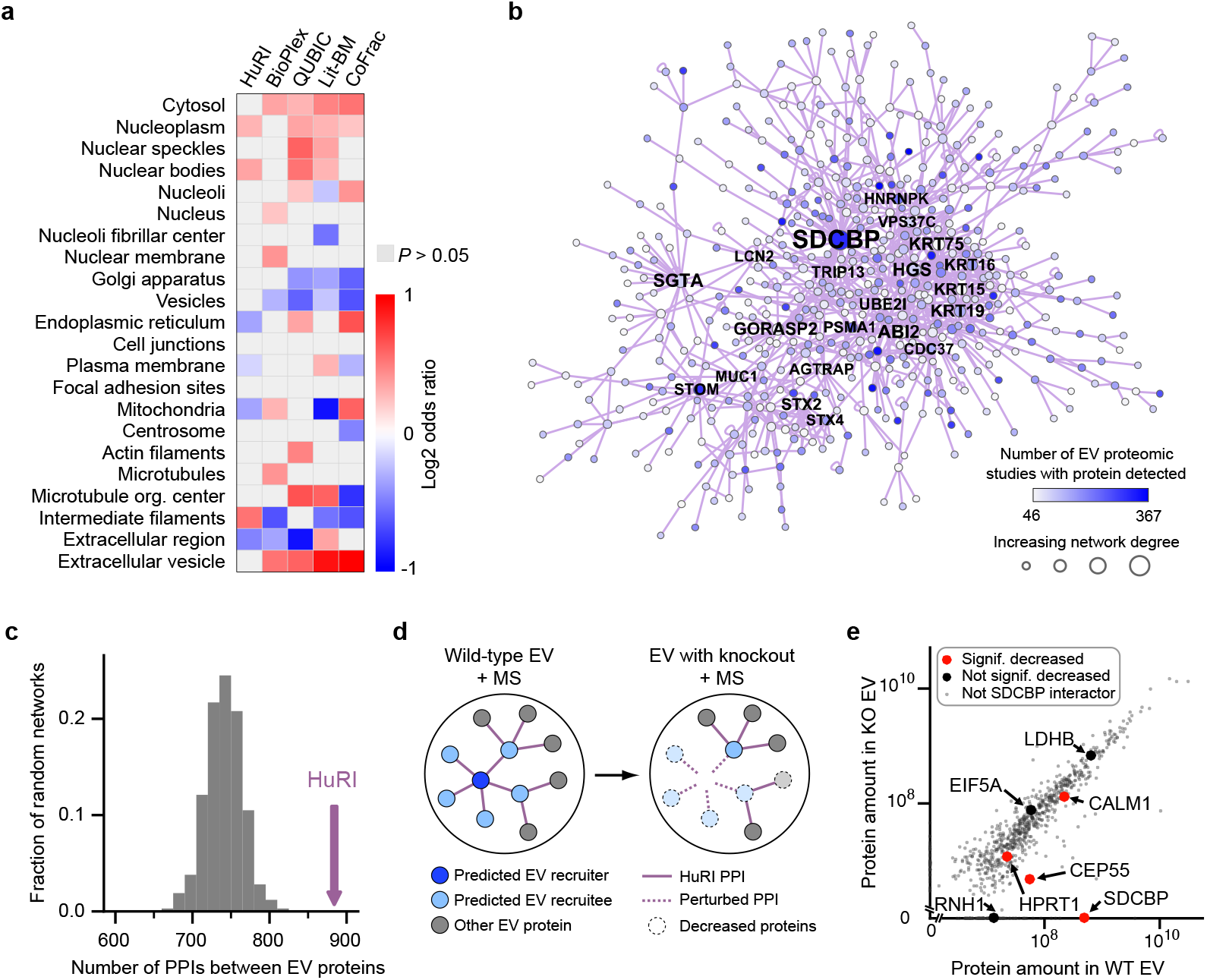
Identification of potential recruiters of proteins into extracellular vesicles. **a**, Odds ratios of proteins in different subcellular compartments and PPI datasets. **b**, The subnetwork of HuRI involving extracellular vesicle (EV) proteins. Names of high degree proteins are shown. **c**, Number of PPIs in HuRI between EV proteins compared to the distribution from randomized networks (grey). **d**, Schematic of experimental design to test EV recruitment function of proteins. MS: Mass Spectrometry. **e**, Protein abundance from EVs for each gene in WT (wild-type) and SDCBP KO (knockout).

One such compartment, extracellular vesicles (EVs), has been intensively studied using proteomics approaches^40^, however, our understanding of the molecular mechanisms that lead to protein recruitment into EVs and subsequent secretion, remains limited. The subnetwork of interactions between EV proteins (Fig. 4b) shows significantly higher connectivity in HuRI than in degree-controlled randomized networks (*P* < 0.001, one-sided empirical test) (Fig. 4c) enabling prediction of EV recruiters using the number of EV interaction partners. Seven of the top 20 most connected proteins in this EV network have established roles in EV biogenesis or cargo recruitment^41,42^ (Fig. 4b). SDCBP (syntenin-1) functions in ESCRT-dependent exosome generation and its knockout shows reduced EV production^43^. SDCBP has 48 PPIs with other EV proteins and is frequently detected in EVs (Fig. 4b), suggesting that it regulates recruitment of interacting proteins to EVs. To test this hypothesis (Fig. 4d), we knocked out SDCBP in the U373vIII cell line (Extended Data Fig. 5a) and found that three of six SDCBP partners detected in the U373vIII EV proteome, CALM1, CEP55 and HPRT1, displayed significantly reduced (*P* < 0.05, one-sided empirical test, fold change < 0.66) (Fig. 4e) protein levels in EVs in the SDCBP knockout line. In contrast, only 15% of the non-interaction partners of SDCBP were reduced (*P* < 0.05, one-sided empirical test) (Extended Data Fig. 5b). Thus, SDCBP may play a role in the recruitment of proteins into EVs, highlighting the potential value of HuRI in studying protein function within specific subcellular contexts.

Despite a significant tendency for interactions in HuRI to link proteins localized to the same compartment (*P* < 0.01, one-sided empirical test) (Fig. 2b), a considerable number of interactions were identified between proteins not reported to co-localize. We find that HuRI PPIs between non-colocalized proteins tended to connect proteins from compartments that significantly overlapped (*P* < 0.001, one-sided empirical test) (Extended Data Fig. 5c-e). This suggests that the lack of co-localization results from incompleteness of the underlying localization annotation and that HuRI could prove useful in predicting additional protein locations and dynamics.

## Principles of tissue-specific cellular function

Despite recent advances in systematic genome-wide identification of tissue-preferentially expressed genes (TiP genes)^2,44^, we lack a concrete understanding of how the surprisingly small set of TiP genes operate together and coordinate their activity with the core housekeeping cellular machinery to mediate tissue-specific functions. Insights can be obtained from investigating the tissue-specific network context of TiP proteins, inferred from integrating protein interactome data with tissue transcriptomes. However, we find that protein complex^7–9^ and literature-curated interactome maps^45^ as well as our previously published binary PPI datasets^10,11^ are strongly depleted for TiP proteins, whereas they are well-represented in HuRI, making it the most suitable interactome map available to study the network context of TiP proteins (Fig. 5a, Extended Data Fig. 6a).

**Fig. 5.**
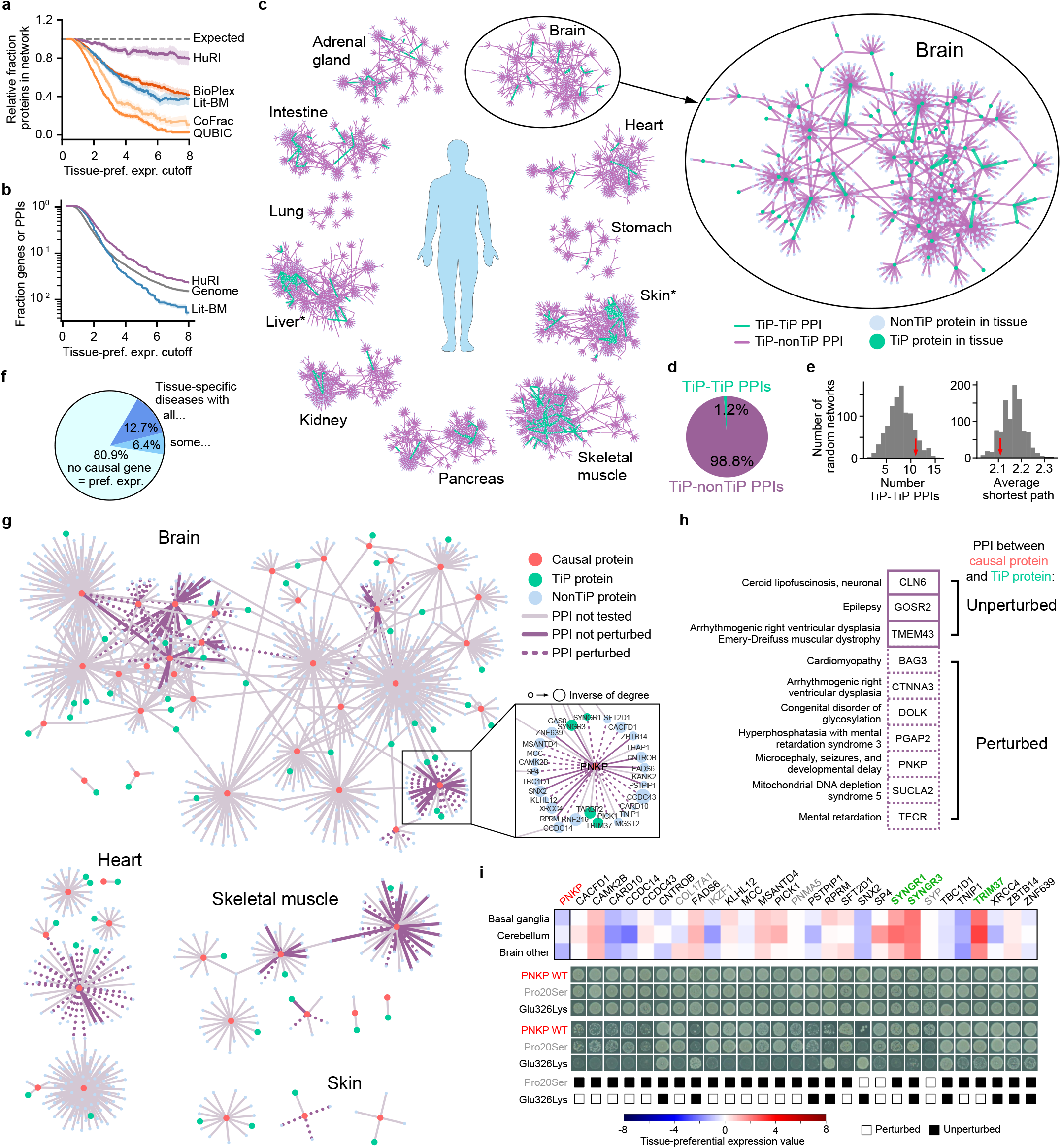
Tissue-specific functions are largely mediated by interactions between TiP proteins and uniformly expressed proteins. **a**, Tissue-preferentially expressed (TiP) protein coverage by PPI networks for increasing levels of tissue-preferential expression. **b**, Fraction of HuRI and Lit-BM that involve TiP proteins compared to fraction of genome that are TiP genes for increasing levels of tissue-preferential expression. **c**, Tissue-preferential subnetworks with enlarged brain sub-network. *: tissues with TiP proteins being significantly close to each other (empirical P < 0.001). **d**, Fraction of PPIs between TiP-TiP proteins in brain. **e**, Empirical test of closeness of TiP proteins in the brain sub-network. **f**, Tissue-specific diseases split by tissue-preferential expression levels of causal genes. **g**, Network neighborhood of uniformly expressed causal proteins of tissue-specific diseases found to interact with TiP proteins in HuRI, indicating PPI perturbation by mutations. **h**, Causal genes split by mutation found to perturb PPI to TiP protein (dashed) or not (solid). **i**, Expression profile of PNKP and interactors in brain tissues and PPI perturbation pattern of disease causing (Glu326Lys) and benign (Pro20Ser) mutation. Yeast growth phenotypes on SC-Leu-Trp (upper) or SC-Leu-Trp-His +3AT media (lower) are shown, green/grey gene symbols: preferentially/not expressed.

System-wide properties of TiP proteins can be determined by assessing their connectivity and centrality in a PPI network compared to “non-TiP” proteins^46^. In HuRI we observe that TiP proteins can engage in as many PPIs and be as central in a PPI network as the more uniformly expressed proteins (degree: Spearman ρ = 0.005, centrality: Spearman ρ = −0.008) (Extended Data Fig. 6b), contrary to previous observations derived from literature-curated PPI networks^47–49^. This result, paired with the fact that PPIs mediated by a TiP protein are effectively also tissue-specific, leads to the finding that the protein interactome as characterized by HuRI is more tissue-specific than the expressed genome (Fig. 5b). This indicates that substantial information on tissue-specific function can only be obtained from the interactome. The opposite is observed for Lit-BM, likely owing to its bias against TiP genes (Fig. 5b).

To investigate the local network neighborhoods of TiP proteins within their respective tissue context, we used HuRI to derive protein interactome maps for 35 tissues^2,50^ (Supplementary Table 14). Each contained an average of 25,000 PPIs that link proteins expressed in that tissue (Extended Data Fig. 6c, d). Within each tissue PPI network, we focused on the interactions involving at least one TiP protein (Fig. 5c). The TiP PPI networks show extensive interactions between TiP proteins and non-TiP proteins, with very few TiP-TiP PPIs dispersed through the network, as exemplified for brain (Fig. 5c, d). Indeed, TiP-TiP PPIs in brain are not enriched, nor is the average shortest path among TiP proteins shorter than in degree-controlled randomized networks (P > 0.05, empirical test) (Fig. 5e). Using either metric, TiP proteins were found to be significantly close to each other in six of 35 tissues, in four of which signals were dominated by clusters of specifically expressed keratins or late-cornified envelope proteins (Extended Data Fig. 6e). Overall, these results provide support for a model in which tissue-specific functions emerge through interactions between TiP proteins and more uniformly expressed members of the basic cellular machinery, presumably modulating and adapting common cellular processes for cellular context-specific needs^51^.

To further investigate functional roles of the identified interactions in HuRI between the basic cellular machinery and preferentially expressed proteins, we selected apoptosis, a biological process with known cell type and developmental stage-specific homeostatic roles^52,53^. We predicted apoptosis-related functions for proteins based on an enrichment of known apoptosis regulators in the protein network neighborhood (Supplementary Table 15). Among the ten most significant predictions were five proteins with demonstrated roles in apoptosis (BCL2L2^54^, BCL2L1^55^, LCN2^56^, BCL2A1^57^ and BCL2L10^58^) supporting the validity of the approach. Among the genes with predicted apoptosis function were C6ORF222, OTUD6A, and NHSL2, three uncharacterized and highly specifically expressed genes (Extended Data Fig. 6f, g). To test the predicted role of the three genes in apoptosis, we assessed their impact on cell viability upon over-expression. Abundance of OTUD6A negatively correlated with time-of-death after addition of TRAIL (TNF-related apoptosis-inducing ligand, *P* = 0.012, two-sided, empirical test, *n* = 40 cells) (Extended Data Fig. 6h), contrary to expression of OTUD6A alone (Extended Data Fig. 6h, Supplementary Table 16). This suggests that OTUD6A participates in the apoptosis pathway but is not itself an inducer of cell death. We found OTUD6A to interact with DYNLL1 and 2 (Extended Data Fig. 6i), two integral members of motor complexes that sequester BCL2L11 and BMF, two “BH3-only” proteins, to the cytoskeleton thereby inhibiting their pro-apoptotic function^59,60^. OTUD6A expression is generally repressed with low expression in eosinophils^61–63^ (Extended Data Fig. 6f) and significant upregulation in response to Decitabine treatment, a drug effective against acute myeloid leukemia^63,64^. Thus, OTUD6A might exert an apoptosis sensitization function via transcriptional activation in a haematopoietic cellular context (Extended Data Fig. 6i).

We were unable to generate a sufficient number of cells expressing C6ORF222 or NHSL2 to perform the cell death assay. However, C6ORF222 contains a BH3 domain^65^ that likely mediates the binding to BCL2L2 and BCL2L1 identified in HuRI (Extended Data Fig. 6i). The interaction between C6ORF222 and the apoptosis regulator MAPK9 identified in HuRI (Extended Data Fig. 6i) was also reported in BioPlex (unpublished released BioPlex data^8^) providing further support for a functional role of C6ORF222 in apoptosis, probably in a digestive tract-specific cellular context (Extended Data Fig. 6g,i). OTUD6A and C6ORF222 represent two examples of specifically expressed genes that might adapt the basic apoptosis machinery to cellular context-specific needs.

## Molecular mechanisms of tissue-specific Mendelian diseases

Many Mendelian diseases display highly tissue-specific phenotypes, which are rarely explained by tissue-specific expression of genes carrying disease-associated mutations^66,67^ (Fig. 5f, Extended Data Fig. 6j). Such mutations have been shown to broadly or specifically affect the formation of PPIs involving the mutated protein^68^. Perturbations of PPIs between uniformly expressed disease-associated proteins and TiP proteins in the corresponding affected tissues have been suggested to underlie the tissue-specific phenotypes of those diseases^67^. Searching the HuRI-derived tissue PPI networks, we find 130 such PPIs involving 63 distinct non-TiP causal proteins and 94 TiP proteins (Fig. 5g). Although we do not observe a significant trend for PPIs between causal proteins and TiP proteins to occur more often than in random networks (Extended Data Fig. 6k), this does not rule out the possibility that perturbations of some of these interactions mediate tissue-specific phenotypes of Mendelian diseases.

To further explore this hypothesis, we experimentally tested whether pathogenic variants associated with the corresponding Mendelian diseases were able to perturb these PPIs. Of ten causal proteins tested, seven showed perturbation of PPIs to preferentially expressed interaction partners in the corresponding “disease tissues” (Fig. 5g-h, Extended Data Fig. 7, Supplementary Table 17). One example is the gene PNKP, mutations in which have been associated with microcephaly, seizures and developmental delay^69^. PNKP, a polynucleotide kinase 3’-phosphatase, is involved in DNA damage repair^69^. The well-established pathogenic mutation (Glu326Lys) affects neither the DNA kinase nor DNA phosphatase activity of PNKP, rendering the mechanism of pathogenicity unclear^70^. We observed that Glu326Lys perturbed PPIs with two partners preferentially expressed in the brain, SYNGR1 and TRIM37, whereas the benign mutation Pro20Ser^71^ did not affect any of these PPIs (Fig. 5i). Interestingly, TRIM37 is known to facilitate DNA repair^72^ suggesting a potential mechanism through which the perturbation of this interaction might affect the brain-specific DNA repair function of PNKP.

In two other cases, CTNNA3 and SUCLA2, the identified TiP interaction partners TRIM54 and ARL6IP1, respectively, themselves cause similar diseases with overlapping symptoms^73,74^, reinforcing the likely pathogenic relevance of the interactions. Overall, this study yields hypotheses of molecular mechanisms for otherwise unexplained tissue-specific phenotypes of seven Mendelian diseases (Fig. 5h) and demonstrates the utility of HuRI as a reference to study biological mechanisms within specific disease contexts.

## Exploration of isoform-specific protein function during development

Transcripts of most human genes undergo alternative splicing, leading in many cases to altered protein sequences. Due to loss or gain of protein interaction-mediating domains or linear motifs, alternative isoforms of the same gene have been found to differ in their interaction profiles^75,76^. Modulation of cellular function by alternative splicing is especially prevalent during developmental processes and in some select adult tissues such as brain or testis^77^, yet most alternative splice products remain uncharacterized. Although we screened only one isoform per gene, HuRI is unique among available human protein interactome maps in providing information about the exact full-length protein sequence of each interaction partner. To aid in the functional characterization of alternative splice products, we aimed at identifying isoforms of a gene with a dominant-negative effect on overall gene function. We combined HuRI with isoform-dependent expression data^78^ to identify genes with isoforms expressed in the same tissue. These genes were further filtered to identify those for which an alternative isoform was predicted to lose some but not all of its interaction-mediating domains^79^, thus likely to lose some but not all of its interaction partners compared to the isoform screened in HuRI^76^ (Fig. 6a).

**Fig. 6.**
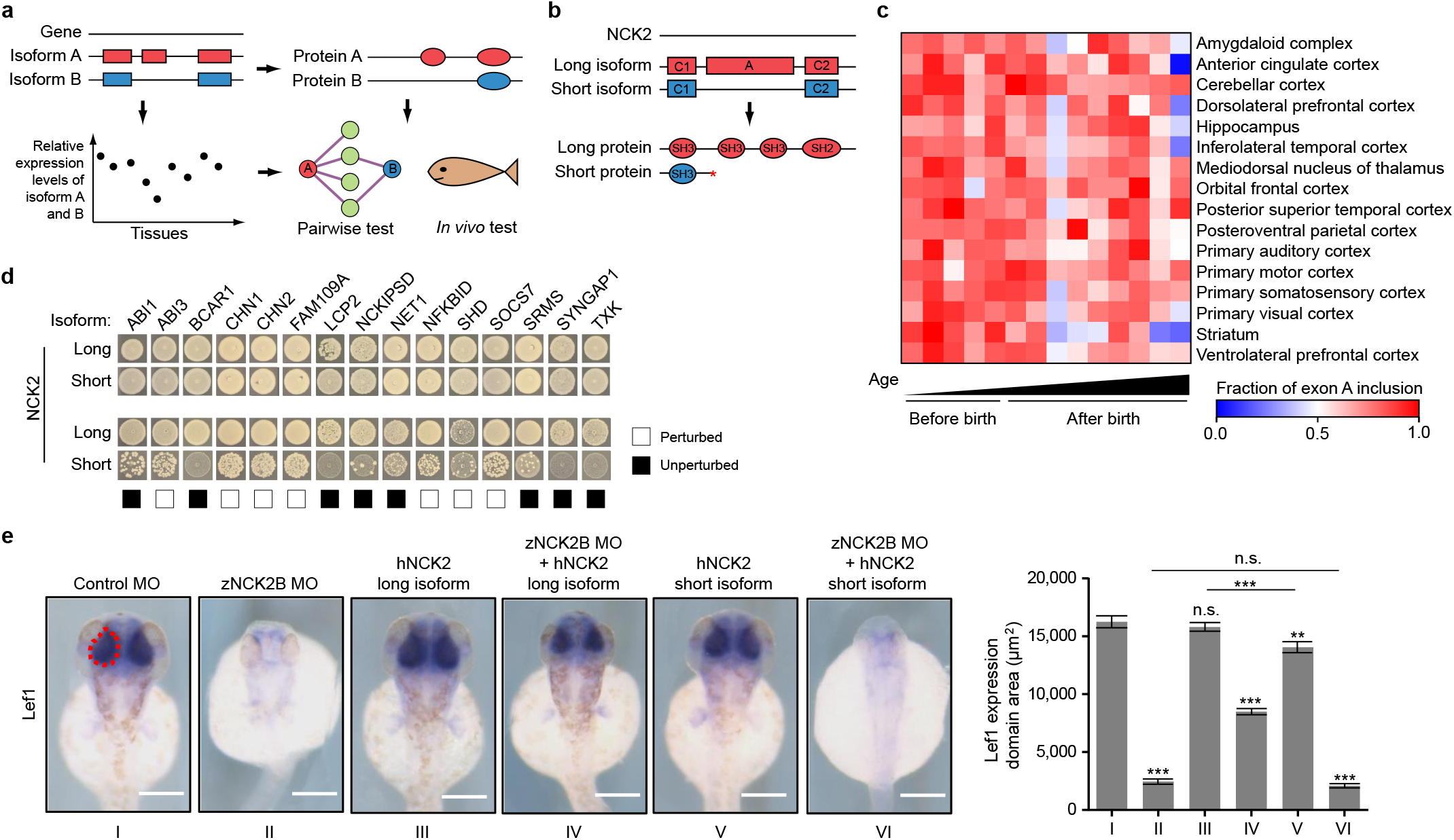
Distinct function of a long and short uncharacterized isoform of NCK2 in brain. **a**, Workflow for predicting and confirming functionally distinct alternative isoforms from the same gene. Rectangles: exons; ovals: protein domains. **b**, Schematic of isoform structure of NCK2. C1, A, C2: constitutive and alternative exons; *: alternative stop codon; rectangles: exons; ovals: protein domains. **c**, Relative NCK2 exon A inclusion in different brain samples. **d**, Pairwise test of long and short isoform of NCK2 with interaction partners in HuRI. Yeast growth phenotypes on SC-Leu-Trp (upper) or SC-Leu-Trp-His+3AT media (lower) are shown. **e**, *In vivo* test of distinct functions of NCK2 isoforms. ***, **, *: P < 0.001, 0.01, 0.05; n.s.: not significant by twosided t-test with n>10; MO: morpholino; zNCK2B: long isoform of NCK2 ortholog in zebrafish. Error bars are standard error.

Of the 192 candidate genes identified (involving known examples of altered alternative isoform function, Supplementary Table 18), we further considered NCK2. NCK2 displayed a well studied principal (long) isoform^80,81^ and an uncharacterized alternative (short) isoform in which three of four predicted interaction-mediating domains are lost (Fig. 6b). Both isoforms are expressed in brain, suggesting specific functional modulation by alternative splicing during brain development (Fig. 6c, Extended Data Fig. 8a, Supplementary Table 19). Known brain-specific functions of NCK2 include synaptic transmission^82^, organization of neuronal circuits^83^, and axon guidance^84,85^. Interestingly, a variant associated with autism was found near a splice site of the alternative exon of NCK2 further suggesting a functional implication of alternative splicing of NCK2 in brain development^86,87^.

Pairwise testing interaction partners of the long isoform of NCK2 with the short isoform confirms the loss of some but not all interaction partners (Fig. 6d) consistent with the retention of one interaction-mediating domain in the short isoform (Fig. 6b). We used zebrafish as a model to test the hypothesized dominant-negative function of the short isoform over the long isoform of NCK2 during brain development (Extended Data Fig. 8b-d). We successfully knocked down both isoforms of zNCK2B in zebrafish using morpholinos (Extended Data Fig. 8e) to measure the impact of loss of zNCK2B on the size of the developing zebrafish brain. While expression of the hNCK2 long isoform in zebrafish under knockdown condition of the endogenous zNCK2B gene partially rescued the phenotype, expression of the short isoform did not (Fig. 6e, Extended Data Fig. 8f, g). Furthermore, expression of the short isoform of hNCK2 in zebrafish brain led to a significant decrease in the size of the developing zebrafish brain, an effect that was not observed with expression of the long isoform of hNCK2 (Fig. 6e). These observations support the predicted dominant negative effect of the short isoform of NCK2 and highlight the power of HuRI to serve as a reference to study isoform-specific protein function in a developmental cellular context.

### Perspectives

By systematically screening about 90% of the protein-coding genome for binary PPIs using a panel of Y2H assays, we generated HuRI, a first reference map of the human protein interactome. Via integration of HuRI with contextual genome, transcriptome and proteome data, we infer cellular context-specific PPI networks that prove to be powerful resources in delineating aspects of cellular context-specific function and organization. Inferred cellular context-specific interactome maps will further gain in accuracy from advances in transcriptomics and proteomics technologies with increasing sensitivities down to the single cell level^4^. Transition from discrete (presence/absence) towards more continuous network models may be achieved by using expression levels to assign weights to interactions^88^, and further improved with large-scale measurements of interaction strengths^9^. Integration of inferred cellular context-specific networks with experimentally-derived cellular context-specific molecular interaction data^89^ will be critical to further refine those models. Future efforts to generate binary protein interactome maps will benefit from development of new PPI assays^16^ capable of identifying PPIs for the proteins still absent from HuRI. New cloning and faster screening technologies^90,91^ are needed to take interactome mapping from testing one isoform per gene to the ensemble of proteoforms generated within the cell. Although multiple challenges remain to be solved for a complete and context-specific map of protein functions, interactions, and higher-level organization, HuRI provides an unbiased genome-scale scaffold with which to coordinate this information as it emerges.

## Acknowledgements

We thank Pablo Porras Millan and the IntAct team for their help in disseminating our PPI data via IntAct, pre- and post-publication. We thank Ulrich Braunschweig, Jonathan Ellis, and Benjamin J. Blencowe for sharing isoform-resolved expression data and discussions on NCK2. We also thank Qian Zhu and Olga G. Troyanskaya as well as Joshua Pan and Cigall Kadoch for sharing co-expression and co-fitness data, respectively. We thank Katharine S. Tuttle for help with graphics. This work was primarily supported by the National Institute of Health (NIH) National Human Genome Research Institute (NHGRI) grant U41HG001715 (M.V., F.P.R., D.E.H., M.A.C., G.D.B, J.T.) with additional support from NIH grants P50HG004233 (M.V. and F.P.R.), U01HL098166 (M.V.), U01HG007690 (M.V.), R01GM109199 (M.A.C.), a Canadian Institute for Health Research (CIHR) Foundation Grant (F.P.R.), the Canada Excellence Research Chairs Program (F.P.R.) and an American Heart Association grant 15CVGPS23430000 (M.V.). D.K. was supported by a Banting Postdoctoral Fellowship through the Natural Sciences and Engineering Research Council (NSERC) of Canada and by the Basic Science Research Program through the National Research Foundation (NRF) of Korea funded by the Ministry of Education (2017R1A6A3A03004385). G.M.S. was supported by NIH Training Grant T32CA009361. M.V. is a Chercheur Qualifié Honoraire from the Fonds de la Recherche Scientifique (FRS-FNRS, Wallonia-Brussels Federation, Belgium).

## Author contributions

The project was conceived and supervised by G.D.B., J.T., D.E.H., M.V., F.P.R. and M.A.C. The Y2H assay versions were developed and benchmarked by K.S., A.D.R., and Q.Z. with help from K.L., B.E.B. and D.B. hORFeome v9.1 was generated by K.L., D.-K.K., K.S., W.B., M.D., D.B., D.M., and T.H. with help from A.G.C., A.Dr., A.M., S.R., Y.S., G.M.S., J.-C.T. and X.Y. The preparation of Y2H destination clones by en masse gateway cloning and yeast transformations were performed by D.-K.K., K.S., A.G.C., J.J.K., R.L., D.M. with help from M.G., D.S., S.S., B.T., C.C., G.H., N.v.L., A.R., and J.W. The Y2H screens were performed and the data analyzed by K.L., K.S., B.E.B., M.D., A.Dr., M.F.H., C.Pol., S.S., B.T., A.T., and T.H. with help from W.B., T.C., B.C., A.De., D.B., S.-F.C., A.M., D.M., J.Ras., A.S.M., Y.S. and Y.W. The validation experiments were performed and the data analyzed by T.C., A.De., I.L., S.G.C., and T.H. with help from K.L., L.L., K.S., D.B., S.D.R., Y.J., Y.K., S.R. and J.T. Sequencing and analysis of the sequencing data was performed by W.B., A.G.C., M.G., N.K., J.K., J.C.M., Y.S. and T.H. with help from K.L., D.-K.K., M.B., C.C., A.G., R.L., A.R., M.T., and J.W. Integrative downstream analyses were performed by K.L., D.-K.K., L.L., F.J.C.-L., I.A.K., and C.Pon. with help from B.C., O.B., G.C., D.D.R., M.D.-F., F.G., G.H., J.N.P., T.R., E.S., E.Y.-L., Y.X., P.A. and J.D.L.R. Follow-up experiments and analysis were performed by K.L., D.-K.K., K.S., R.B., D. C., S.D., A.De. and A.Y. with help from L.L., T.C., C.B.-C., G.C., C.D.A., H.E., L.G., E.H., S.La. and R.J.W. supervised by S.G., J.Rak. and V.T. The webportal was developed by M.W.M. with help from K.L., M.H., T.H., M.A.C. and G.D.B. The paper was written by K.L., D.-K.K., L.L., K.S., D.E.H., M.V., F.P.R., and M.A.C. with help from F.J.C.-L., A.G.C., G.C., S.G., I.A.K., T.H, and A.Y. Authors other than co-first and co-corresponding are listed alphabetically and contributed equally within their group.

## METHODS

### Construction of Reference ORFeome and definition of screening space (space III)

#### Selection of existing clones

We supplemented our human ORFeome collection hORFeome version 7.1 (v7.1) (http://horfdb.dfci.harvard.edu/hv7)^92^ with clones from additional genes from the ORFeome Collaboration^93^ (http://www.orfeomecollaboration.org), DNASU plasmid repository^94^ (https://dnasu.org) and other collaborators. All clones are in Gateway compatible entry vectors with spectinomycin or kanamycin resistance markers, as appropriate. While native stop codons of most clones have been removed, 617 clones contain native stop codons.

To select a single Open Reading Frame (ORF) for genes with multiple ORFs available, for each gene, we aligned the sequences of the corresponding ORFs to human genome GRCh37 using BLAT^95^ (v36×1) and chose the longest ORF with more than 95% of its sequence aligned to the genome. If no such ORF was available, we chose the ORF with the highest percentage of alignment to the genome.

#### Cloning of new ORFs

After collecting available ORFs from different resources, about ~3,000 human protein-coding genes remained uncovered (no ORF available). To obtain ORF clones for these missing genes, we attempted RT-PCR on a pool of cDNA libraries from brain, heart, and liver ordered from Biochain (Human Adult Normal Tissue: Brain, catalog number C1234035, lot number C203351; Human Adult Normal Tissue: Heart, catalog number C1234122, lot number B901100; Human Adult Normal Tissue: Liver, catalog number C1234149, lot number C203352). We designed primers for all the missing genes for which we could find coding sequences from RefSeq^96^ (https://www.ncbi.nlm.nih.gov/refseq; downloaded April 13th, 2015). In total, we attempted 2,257 primer pairs and successfully cloned 481 ORFs into pDONR223 vector, all without native stops. The sequences of the clones were verified using Illumina sequencing. We named the combined collection hORFeome v9.1.

#### Definition of the human protein-coding genome

GENCODE^97^ annotation (ftp://ftp.sanger.ac.uk/pub/gencode/Gencode_human/release_27) and DNA and protein sequences were filtered for information on transcript entries. After removing PseudoAutosomal Regions on the Y chromosome (PAR) genes, all genes of gene_type “protein_coding” with annotated transcripts yielded the 19,818 protein-coding genes used in this study. External datasets were mapped via gene or protein IDentifiers (IDs) to the Ensembl gene ID space, and genes, proteins, or transcripts that did not map were removed.

#### Mapping of ORFs to GENCODE release 27

To assign Ensembl gene IDs to ORFs in the hORFeome v9.1 collection, we aligned ORFs to protein-coding transcripts in GENCODE^97^ and determined the best match for each ORF-transcript pair. Briefly, ORF protein sequences were aligned to all protein sequences of protein-coding transcripts using BLASTP^98^ (NCBI BLAST v2.2.30) with default parameters. Alignments were further refined using MUSCLE^99^ (v3.8.31; default parameters). For each Ensembl gene, we kept only alignments with identity ≥ 95%, ORF coverage ≥ 50%, transcript coverage either ≥ 50% or with at least two exons covered. The best match was determined based on the combination of ORF coverage, transcript coverage and alignment identity.

Each ORF could generally be assigned to only one Ensembl gene, but where alignments were fully identical among different genes (e.g. histones), the ORF was assigned to multiple genes. Finally, we removed 33 ORFs containing known disease mutations based on HGMD^100^ v2016 annotation. Using this pipeline, hORFeome v9.1 was matched to 17,408 protein-coding genes (Supplementary Table 20).

### Generation and benchmarking of Yeast Two-Hybrid (Y2H) assay versions

#### Vector design

To generate pDest-AD-AR67 [Activation Domain (AD) at C-terminus (C-term) with yeast centromere (CEN)], a fragment encoding an *ADH1* promoter, Gateway recombination cassette, and C-terminal AD was PCR amplified from pGADCg^101^ using forward primer AP36 (5’ GAAGGCTTTAATTTGCAAAGCTCGGGATCCGGGCCCCCCCTCGAGATCCGcatctattgaagtaataataggcgcatg 3’) and reverse primer AP37 (5’ CAACCTTGATTGGAGACTTGACCAAACCTCTGGCGAAGAAGTCCAAAGCTctgaataagccctcgtaatatattttcatg 3’) and cloned into EcoRI (New England Biolabs, NEB) and SalI (NEB) digested pDEST-AD via homologous recombination gap repair in yeast. To generate pDest-AD-AR68 (AD at Cterm with yeast 2μ), the same fragment encoding the ***ADH1*** promoter, Gateway recombination cassette, and C-terminal AD was instead PCR amplified from pGADCg using forward primer AP36 and reverse primer AP38 (5’ GCTGCATGTGTCAGAGGTTTTCACCGTCATCACCGAAACGCGCGAGGCAGcatctattgaagtaataataggcgcatg 3’) and cloned into NotI- and XmaI-digested (NEB) pDest-AD-QZ213 via homologous recombination in yeast. See Supplementary Table 21 for details on vector design and Y2H assay versions.

#### Benchmarking pairwise test performance

Assay versions were benchmarked by Y2H using a positive reference set of 92 well-documented interacting human protein pairs (Positive Reference Set; PRS v1) and a set of 92 random human protein pairs (Random Reference Set; RRS v1), as previously described^14^. Briefly, haploid yeast strains Y8800 and Y8930 carrying plasmids expressing AD and DB reference set fusions, respectively, were mated overnight and diploid selection was performed. Diploids were spotted onto Synthetic Complete media without Leucine, Tryptophan and Histidine with 1 mM 3-Amino-1,2,4-triazole (SC-Leu-Trp-His+1 mM 3AT for assay versions 1 & 2) or SC-Leu-Trp-His+10 mM 3AT (for assay version 3) solid media, incubated at 30°C for 4 days, and interactions were scored based on the strength of *GAL1::HIS3@LYS2* reporter activation. Pairs that displayed AD-independent *GAL1::HIS3@LYS2* autoactivation were designated as negatives.

#### Benchmarking screening capabilities of Y2H assay versions on a test space

For each assay version we performed several screens of a test space^10^ covering ~1% of the total space (~2,000 DB’s against ~2,000 AD’s) to adjust the protocol for high-throughput screening. Based on that, we were able to calculate how many screens per assay version are needed to reach saturation and how many new pairs are found by screening a space several times. For Y2H assay version 1, 2 and 3, we performed 12, 3, and 6 screens, respectively. The screens were performed and validated as described below in the Y2H screening, retest, and validation sections.

### Subcloning into Y2H vectors

#### Preparation of Y2H destination clones by 1-to-1 Gateway cloning

ORFs from the hORFeome v9.1 collection were transferred by Gateway recombinational cloning (Invitrogen) into Y2H destination vectors pDEST-DB and pDEST-AD-CYH2 to generate DB and AD-hybrid proteins (DB-ORF and AD-ORF, respectively), as described previously^102^. In addition, all ORFs were transferred into pDest-AD-QZ213 for assay version 2 and all ORFs without a stop codon into pDest-AD-AR68 for assay version 3.

#### Preparation of Y2H destination clones by en masse Gateway cloning

To increase the throughput and efficiency of the cloning, we transferred the majority of the clones into the pDest-AD-AR68 vector *en masse*. To enable future use in the barcode fusion genetics Y2H system^90^. As described previously^90,91^, we generated randomly barcoded Y2H destination plasmid pools and carried out *en masse* Gateway LR reactions, bacterial transformations, colony pickings, and sequencing to identify ORFs and barcodes. After obtaining the raw sequencing reads, we ran Illumina bcl2fastq (v2.20 with options “--no-lane-splitting -r 3 -p 10 -w 3”) to demultiplex all the reads into different plates according to the i5/i7 index sequences. Extracting well-tag from each read, identified by locating conserved flank sequences, allowed assignment of reads from each plate to the corresponding well of origin. Full length (24-26 nt) barcodes within one base-pair mismatch were merged to identify the dominant barcode for each well. ORF reads in each well were aligned to reference ORF sequences via Bowtie 2^103^ (v2.2.3). For wells containing more than one clone, we filtered all the ORF and barcode pairs found in each well by calculating the percentage of reads aligned to each ORF. Only dominant barcoded-ORFs were selected, as defined by the ORF with the highest fraction of reads (at least, 20 reads and 20%) and ≥5X more reads than those for the second dominant ORF. About 1% of clones were validated by Sanger sequencing with ≥95% validation rate.

### Primary screening

#### Yeast strains and transformation

Competent yeast strains Y8800, mating type MATa, and Y8930, mating type MATa, both harboring the genotype leu2-3,112 trp1-901 his3Δ200 ura3-52 gal4Δ gal80Δ GAL2::ADE2 GAL1::HIS3@LYS2 GAL7::lacZ@MET2cyh2R were transformed with individual AD-ORF and DB-ORF constructs respectively, and plated onto SC-Trp or SC-Leu to select for AD-ORF or DB-ORF plasmids^102^.

#### Auto-activator identification and removal

Prior to Y2H screening, haploid DB-ORF yeast strains were tested for auto-activation of the GAL1::HIS3 reporter gene in the absence of any AD-ORF plasmid. Individual DB-ORF yeast strains showing growth in a spotting assay on SC-His-Leu+3AT media were considered autoactivators and removed from the collection of strains to be screened.

#### Y2H screening

Fresh overnight cultures of individual Y8930:DB-ORF yeast strains (bait) were mated against Y8800:AD-ORF libraries containing ~1,000 different Y8800:AD-ORF yeast strains (prey). After overnight growth at 30°C in liquid rich medium (YEPD), mated yeast cells were transferred into liquid SC-Leu-Trp media to select for diploids. After overnight incubation at 30°C diploid yeast cells were spotted onto SC-Leu-Trp-His+3AT solid media to select for activation of the *GAL1::HIS3* reporter gene. In parallel, diploid yeast cells were transferred onto SC-Leu-His+3AT solid media supplemented with 1 or 10 mg/l CHX for assay version 1 or 2 and 3, respectively. All AD-ORF plasmids carry the counter-selectable marker CYH2, which allows selection on CHX-containing medium of yeast cells that do not contain any AD-ORF plasmid in order to identify spontaneous DB-ORF auto-activators^102^. After 72h incubation at 30°C, yeast that grew on SC-Leu-Trp-His+3AT media but not on SC-Leu-His+3AT+CHX media were picked into SC-Leu-Trp overnight and then processed to determine the identity of the respective bait and prey proteins.

#### Yeast colony PCR and sequencing

Because each DB-ORF yeast strain was mated against a library of ~1,000 AD-ORF yeast strains in the first-pass screens, one bait protein could interact with more than one prey proteins per mini-library. To account for this, we picked between three and five colonies (primary positives) from each growth spot on SC-Leu-Trp-His+3AT media to 96-well plates. Primary positive colonies were processed as described^102^ to generate lysates for PCR. Three microliters of diluted lysate was used as a template for PCR amplification to generate DB-ORF and ADORF PCR amplicons.

To cost-effectively identify both bait and prey proteins for hundreds of thousands positive colonies, we developed a method called SWIM-seq (Shared-well Interaction Mapping by sequencing). Using SWIM-seq, we took advantage of next generation sequencing technology and pooled DB-ORF and AD-ORF PCR products that contain well-specific and plate-specific nucleotide sequences from tens of thousands colonies together in one sequencing run. Briefly, PCR was performed using AD and DB well index primers together with universal primers (see table below). All PCR was done using Platinum Taq (Life Technologies). After PCR, the products from each PCR plate were pooled into one single well in a 96-well plate (SWIM plate). The SWIM plates were purified (Qiagen, PCR Purification Kit) and processed to make an Illumina sequencing library, during which Illumina adapter sequences, i7 and i5, were incorporated as plate indexes. The library was then paired-end-sequenced using an Illumina platform (MiSeq or NextSeq 500). See Supplementary Table 22 for the primers used for the different Y2H assay versions.

#### Identification of pairs of AD-ORF and DB-ORF

We developed a computational pipeline to process demultiplexed paired-end reads and identify the matching ORF pairs corresponding to Y2H-positive colonies. Paired-end reads are in a fastq format, with one read, R1, containing a part of the ORF sequence and the other paired read, R2, containing the well index. Briefly, we used Bowtie 2^103^ (v2.2.3) to align all R1 reads to reference sequences and extracted the well-identifying indices from the R2 reads. AD-ORFs and DB-ORFs that shared the same well indices were paired together. Pairs identified from the primary screen were calls FiPPs (First Pass Pairs). To identify likely true interacting pairs, we developed a “SWIM score” S that takes into account the AD and DB reads in each well, total reads returned from the sequencing run, and other factors.

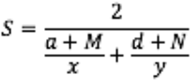

where *x* and *y* are read counts of an AD-ORF and DB-ORF in a given well respectively, *a* and *d* are total read counts of all aligned AD-ORF and DB-ORF in that well, and *M* and *N* are pseudocounts for AD and DB respectively, which were constant for each sequencing batch but varied for different batches.

We then selected FiPPs for pairwise test using a cutoff that balances the risk of testing too many FiPPs that are not protein-protein interactions (PPIs) versus not testing too many FiPPs that are actual PPIs. The cutoff varied for different screens and sequencing runs to accommodate slight variations in the screening and sequencing protocol.

### Pairwise test

#### Design of pairwise test and scoring

No FiPP is considered a PPI and released before it has been verified in a pairwise test using Y2H. For all FiPPs, each protein was picked from the stock collection and mated with its partner. This experiment was done in quadruplicate, with yeast spotted on both SC-Leu-Trp-His+3AT plates and SC-Leu-His+3AT+CHX to test for spontaneous auto-activation. As positive and negative controls in each test batch, 92 pairs from PRS v1, 250 randomly selected Lit-BM pairs (literature binary multiple pairs), and 250 pairs of proteins randomly selected from the search space were included. Experiments were considered successful if at most 1% of the random protein pairs and 10% or more of the Lit-BM or PRS v1 pairs scored positive. Positive and negative controls were combined in the same batches with FiPPs and scored blindly. To ensure the mating of the corresponding plates with each other (AD-X with DB-Y) and identify any potential plate rotations or swaps, each retest plate has a unique pattern of empty wells. We developed a pipeline to semi-automate the scoring of growing yeast in 384 or 96 format. Growing yeast colonies were first scored automatically by software that uses the python library Colonyzer^104^ to process the images. The scores were then manually checked prior to saving and pixel intensities were used to estimate the strength of growth. Both retest plates and CHX control plates were independently scored and scores combined to produce a final score of each well. A pair was scored invalid (NA) for 96-well format if the well was unscorable (contaminated, not spotted, etc.); or, if in 384 format, if at least 2 of the 4 quadruplicates were unscorable on the retest plate; or, if the corresponding well on the CHX plate was unscorable. If a pair was not scored as NA but grew at least as strongly on the CHX plate compared to the growth on the retest plate (average of quadruplicates), then it was scored as a spontaneous auto-activator. If a pair was neither NA nor an auto-activator and showed growth in at least 3 of 4 replicates on the retest plate, then it was scored as positive; otherwise, negative.

#### Treatment of pairs scored as spontaneous auto-activators

Because validation data has suggested that the CHX control is over-sensitive for 2μ vectors and removes many actual PPIs, we performed one more pairwise test for pairs that scored as spontaneous auto-activators in the pairwise test in assay versions 2 and 3. For these retests, the DB-ORF was separately mated with an “AD-nul” plasmid without any ORF in the cloning site. If a yeast colony grew more strongly when mated to the corresponding AD-ORF than the AD-null strain, we “rescued” this PPI and added it to the dataset after sequence confirmation.

#### Sequence confirmation of positive pairs from pairwise test

To guide the picking of all positive pairs for sequence confirmation, a “picking map” with all positive pairs was generated by the scoring software and printed. Picked pairs were lysed and SWIM-PCR was performed as described in the above section. Illumina reads were processed via the computational pipeline described above. Only pairs identified from the pipeline that matched the tested ORF pairs previously scored as positive were considered PPIs.

### Validation of screens with orthogonal PPI assays

#### Design of validation experiments

To validate each of the nine screens of Space III, we randomly selected ~340 positives from each screen and tested them in several batches using two assays, MAPPIT (MAmmalian Protein-Protein Interaction Trap) and GPCA (*Gaussia princeps* luciferase protein complementation). For each batch, we included ~200 randomly selected PPIs from Lit-BM as PRS and ~400 pairs of proteins randomly selected from the search space as RRS. For each pair, a random bait-prey configuration was assigned. All tested pairs and control pairs were combined in batches and tested blindly.

For the test space validation, unlike in the main screen where we chose PPIs for each screen, we randomly chose PPIs from all positive pairs together regardless of which screens they were from. All Lit-BM PPIs (~200) and ~400 random pairs of proteins from within the tested space were treated as a positive reference set and a random reference set respectively.

#### MAPPIT assay

MAPPIT experiments were performed as previously described^10^. Briefly, HEK293T cells were grown in 384-well plates and co-transfected with a luciferase reporter and plasmids for both bait and prey fusion proteins. Twenty-four hours post-transfection, cells were either stimulated with ligand (erythropoietin) or left untreated, then incubated for an additional 24 hours before luciferase activity was measured in duplicate. The MAPPIT validation experiment was deemed valid if both bait and prey were successfully cloned into expression vectors and bait expression was detected. “Fold-induction” values (signal from stimulated cells divided by signal from unstimulated cells) were calculated for each tested pair, and two negative controls (no bait with prey and bait with no prey). Each tested pair was assigned a quantitative score: the fold-induction value of the pair divided by the maximum of the fold-induction value of the two negative controls.

#### GPCA assay

As an orthogonal validation assay, GPCA experiments were performed as described elsewhere^16^. Briefly, GPCA N1 and N2 vectors are based on two fragments of the *Gaussia princeps* luciferase with humanized codon usage (herein referred as GLuc)^19^. Both GLuc fragments were linked to the N-terminus of the tested proteins via a flexible linker polypeptide of 20 amino acid residues, including the Gateway recombinational cloning sites. To normalize mRNA translation initiation, a consensus Kozak translation start sequence was added. Both constructs were carried by the same CytoMegaloVirus (CMV)-driven mammalian expression vector (pCI-neo derived, Promega) and were maintained at high copy number via the presence of the SV40 replication origin.

HEK293T cells were seeded at 6×10^4^ cells per well in 96-well, flat-bottom, cell culture microplates (Greiner Bio-One), and cultured in Dulbecco’s Modified Eagle’s Medium (DMEM) supplemented with 10% fetal calf serum at 37°C and 5% CO_2_. 100 ng of purified plasmid DNA for each protein of a pair was transfected into HEK293T cells in 96-well, flat bottom, cell culture plates (Greiner Bio-One) supplemented with 10% Fetal Bovine Serum (FBS) in DMEM using PolyEthylenImine (PEI)^19^. The DNA/PEI ratio (mass:mass) was 1:3. Cell culture medium was removed 24 h after DNA transfection, and cells were gently washed with 150 μL pre-warmed 1x PBS (phosphate buffered saline). 40 μL of lysis buffer was added to each well, and cell lysis was performed under vigorous plate shaking for 20 min at 900 rpm. Luminescence was measured by auto-injecting 50 μL Renilla luciferase substrate (Renilla Luciferase Assay system, Promega) in each well and integrating light output for 4 s using a TriStar luminometer (Berthold) to obtain quantitative scores. GPCA has not been performed on PPIs of screens 7, 8, and 9 generated with Y2H assay version 3.

#### Processing of data and calculation of validation rates

In MAPPIT and GPCA assays, if a pair is positive or negative was determined by thresholds set at the 99th percentile of the RRS scores (equivalent to a 1% false discovery rate). This was determined separately for each experimental batch and calculated using the quantile function in the Python pandas library. Pairs without valid quantitative scores were dropped, and recovery rates were calculated as the number of positive pairs over the sum of the positive and negative pairs. The error bars on the recovery rates were calculated using a Bayesian model (a binomial likelihood with a uniform prior), taking the central 68.27% interval of *Beta* (*p* + 1, *n* + 1), where *p* and *n* are the number of pairs testing positive and negative, respectively.

#### Testing of PPIs in MAPPIT that were found in multiple screens and analysis of data

Randomly sampled PPIs were selected for validation assessment from individual screens and from additional randomly-sampled pairs from subsets of PPIs that were defined by the number of screens in which the PPI was detected. Every pair in HuRI that was also in any of the literature curated datasets [Lit-BM / Lit-BS (literature binary singleton pairs) / Lit-NB (literature not binary)] was also selected for validation assessment.

### Processing of external transcriptome datasets: GTEx, FANTOM and Human Protein Atlas

Genotype-Tissue Expression project (GTEx)^2^ v6 transcriptome data previously processed with the R YARN package^50^ was downloaded from http://networkmedicine.org:3838/gtex_data/gtex_portal_normalized.rds on March 15th, 2016 and processed with R v3.5.1 to extract normalized log read count data for every gene and tissue sample. The median expression of every gene across all samples from a given subtype (tissue or cell line) was calculated and considered as the expression level of the genes in those tissues. A median expression cutoff of >5 was applied to consider a gene being expressed in a given tissue. Cell line samples were excluded and genes were restricted to the set of protein-coding genes (see above). The YARN package collapses the 16 brain subregions into three brain tissues, basal ganglia, cerebellum, and other, which have been used in this study (Supplementary Table 23).

Transcripts Per Kilobase Million (TPM) from Cap Analysis Gene Expression (CAGE) peak data was extracted from the hg19.cage_peak_phase1and2combined_tpm.osc.txt file (http://fantom.gsc.riken.jp/5/datafiles/latest/extra/CAGE_peaks/; version November 11th, 2014). Entrez gene IDs were mapped to Ensembl gene IDs and the dataset was restricted to protein-coding genes. For each gene from FANTOM^3^, we computed the mean of cage peak TPM values of all samples associated to any FANTOM primary cell category. Samples were mapped to primary cells using FANTOM SSTAR (http://fantom.gsc.riken.jp/5/sstar/). Primary cell categories that were considered to be biologically too similar and of close TPM expression were merged. TPM values were log2 transformed and a gene was considered expressed if its value > 0.

RNA-seq data from 64 cell lines processed at the gene level with TPM values downloaded on December 1st, 2017 from the Human Protein Atlas^13^ (https://proteinatlas.org). Genes with TPM ≥ 1 were considered expressed.

### Computing increase of HuRI with more assay versions and screens

To evaluate the number of PPIs and proteins observed as a function of the number and assay version of screens used, the mean of the cumulative number of PPIs and proteins across all matching screen combinations was used. For example, for five screens with three being assay version 1 and two being assay version 2, the mean number of unique PPIs was calculated for all combinations of any 3 of the 12 assay version 1 screens and any 2 of the 3 assay version 2 screens.

### Integrative analyses with protein structural information

#### Analysis of distances between protein termini and interaction interfaces

We retrieved experimental structures, or either complete or domain-based models (using Interactome3D^21^ version 2018_04) involving any two proteins that have at least one interaction in HuRI. For each subunit in a complex structure, we defined its interaction interface as the residues for which the Accessible Surface Area (ASA) changed more than 1 Å^2^ between the bound and unbound state. The center of this interaction interface was determined as the Cα atom being closest to the center of coordinates of the interaction interface. Next, we calculated the distance from the N- and C-terminal Cα atoms to the center of the interaction interface. Importantly, we only considered protein structures with complete N- or C-termini depending on the analysis performed. In complexes with truncated tails, we searched for monomer structures with a minimum of 50% coverage and complete terminal tails. In those cases, we superimposed the monomers onto the respective complex subunits and calculated the distance from the relevant terminus to the interaction interface center. Finally, we grouped these interactions based on the Y2H assay version(s) in which they were detected and performed the analysis separately for DB and AD fusions.

#### Analysis of the fraction of PPIs in dataset that are direct

We queried Interactome3D^21^ (version 2018_04) for complexes involving three or more proteins with experimental structure available. For all combinations of protein pairs within a complex, Interactome3D calculated the number of residue-residue contacts by accounting for hydrogen bonds, van der Waals interactions, and salt and disulfide bridges. We defined protein pairs with five or more contacts as direct, and remaining pairs as indirect. Separately for each given PPI dataset, the fraction of direct PPIs was calculated as the number of direct PPIs divided by the number of direct and indirect PPIs.

#### Analysis of correlation between number of screens and interaction strength

We downloaded from Interactome3D^21^ (version 2018_04) the experimental structures involving any two proteins in HuRI (Human Reference Interactome). For each complex structure, we calculated the interaction interface area by subtracting the ASA in the bound state from the total ASA of the unbound proteins and by dividing the result by two. Additionally, we calculated the number of residue-residue contacts as explained above. Finally, we grouped the protein pairs by the number of HuRI screens in which an interaction was detected.

### Retrieval and processing of other interactome datasets

#### Construction of Lit-BM

Literature curated datasets were derived from the Mentha resource^45^, which aggregates literature-curated PPIs from five source databases: MINT^105^, IntAct^106^, DIP^107^, MatrixDB^108^ and BioGRID^109^. The data downloaded from Mentha was dated August 28th 2017. Data was filtered to have valid IDs for the UniProt accession numbers, Pubmed IDs and PSI-MI terms. Each piece of evidence for a protein pair must consist of a Pubmed ID and an interaction detection method code in the PSI-MI controlled vocabulary^110^. Duplicated evidence arises in cases where different source databases curate the same paper. We merged duplicated entries for each pair, as detected by multiple pieces of evidence with the same Pubmed ID and experimental interaction detection codes which are either the same or have an ancestor-descendent relationship in the PSI-MI ontology. (In the latter case, the more specific descendent term was assigned to the merged evidence.) In order to select the subset of PPIs corresponding to binary interactions (as opposed to co-complex associations) we developed a manual classification of the PSI-MI interaction detection method^110^ terms, which define the experimental technique used. Our classification has been updated since previous versions, in order to cover new experimental methods which have been added to the controlled vocabulary in the intervening time. The methods are classified into three categories; ‘invalid’, ‘binary’ and ‘indirect’. Where ‘invalid’ corresponds to terms that are not considered valid experimental protein-protein interaction detection methods, ‘binary’ to terms that detect binary protein-protein interactions and ‘indirect’ to terms that detect potentially indirect interactions. An example term in the first category is “colocalization”. All PPI evidences annotated with “invalid” terms are removed and not considered. The other two categories are used to divide the protein pairs in the literature-curated dataset into three categories, as follows. If a pair has no evidence that correspond to a binary method it is classified as Lit-NB, if a pair has only one piece of evidence with a binary method it is classified as Lit-BS and if a pair has multiple pieces of evidence, with at least one corresponding to a binary method then it is annotated as Lit-BM. The resulting number of PPIs in each category is shown in (Extended Data Fig. 9a).

The interactions that make up the literature come from experiments that span a broad range of different sizes. It is possible that there could be a difference in average quality between experiments that report just a single or small number of PPIs and those that use high-throughput techniques to identify large numbers of PPIs. In order to investigate this we divided the Lit-BM and Lit-BS into High-Throughput (HT) and Low-Throughput (LT) subsets, based on the size of the smallest experiment, that provided a piece of evidence for that pair using a binary experimental method. If 50 or more PPIs were reported in that experiment, it was classified as HT and otherwise was classified as LT. Random samples of these subsets were pairwise tested using Y2H (assay version 1) and MAPPIT (Extended Data Fig. 9b, c, Supplementary Table 24, 25). We observed no significant difference between the validation rates of the HT and LT subsets and so we made the decision not to implement any cutoff in experiment size when defining the literature dataset.

The main differences from the previous literature curated dataset Lit-BM-13^10^ are the databases used. Random samples of Lit-BM-17 pairs show similar recovery rates to random samples of Lit-BM-13 pairs in Y2H and MAPPIT (Extended Data Fig. 9d, e). PPIs from Lit-BM-13 have been used as positive controls in experiments, PPIs from Lit-BM-17 (Supplementary Table 26) have been used in all computational analyses.

#### Processing of co-complex interactome maps BioPlex, QUBIC, CoFrac

BioPlex^8^ v2.0 data was downloaded from http://bioplex.hms.harvard.edu on August 1st, 2018, and protein IDs were mapped from UniProt to Ensembl gene IDs. QUBIC was downloaded from the supplementary table of the previous study^9^ and protein IDs were mapped from UniProt to Ensembl gene IDs. CoFrac^7^ was processed by downloading the data from http://human.med.utoronto.ca on August 1st, 2018 and mapping from UniProt to Ensembl gene IDs. All interatome maps were then filtered to contain interactions only involving Ensembl gene IDs corresponding to protein-coding genes.

#### Processing of previously generated binary interactome maps at CCSB

To update the previous CCSB (the Center for Cancer Systems Biology) binary interactome maps and convert them into Ensembl gene ID pairs, for each map, we retrieved the original ORF based IDs stored in our database and mapped the ORF IDs to Ensembl gene IDs as described above. PPIs involving ORFs that remained unmapped or were discarded or mapped to non-protein-coding genes were removed.

### Analysis of HuRI PPIs linking proteins within versus between protein complexes

CORUM^22^ complexes were downloaded from http://mips.helmholtz-muenchen.de on August 1st 2018 and gene IDs were mapped from UniProt to Ensembl. Complexes with less than three subunits were removed as well as complexes that overlapped 90% or more with another complex (the smaller complex of both was removed). HuRI PPIs were filtered to remove homodimers and were restricted to those linking proteins annotated to be in at least one CORUM complex (restriction to common space between HuRI and CORUM). The fraction of these HuRI PPIs that link a pair of interacting proteins that were observed to be at least once part of the same CORUM complex was determined and these PPIs were further split into PPIs found in only one of the 9 HuRI screens versus having been found in at least 2 screens. The fractions plotted were 137/1042 and 232/775 for one screens and multiple screens, respectively. CORUM complex membership was used to identify all interactions between them from BioPlex^8^ or HuRI. py2cytoscape^111^ was used to generate network models of each complex.

### Functional enrichment analyses of HuRI and the Profile Similarity Network (PSN) of HuRI

#### Construction of PSN and randomizations

Jaccard similarity (number of shared interaction partners divided by union of interaction partners) was calculated for every pair of proteins of degree ≥ 2 in a given network. Only pairs of proteins that shared at least one interaction partner were considered in any further analysis using the PSN. Random PSNs were generated in the same way from degree-controlled randomized HuRI networks that generated using the degree_sequence() function in the Python igraph^112^ library v0.7.1.

#### Calculation of pairwise sequence identity and correlation with Jaccard similarity

To calculate the sequence identity between a pair of proteins in HuRI, we aligned the two corresponding ORF protein sequences using MUSCLE^99^ (v3.8.31) with default parameters. We then parsed the MUSCLE alignment file and calculated the sequence identity as the number of matched amino acid positions divided by the length of the alignment. To assess whether higher Jaccard similarity between two proteins in HuRI correlates with higher overall sequence similarity between the two proteins as an indication for similar interaction interfaces while controlling for degree biases, we computed for increasing sequence identity cutoffs the sum of Jaccard similarities of all edges in the PSN that link two proteins with a sequence identity of or above that cutoff. This sum was divided by the mean of the sums calculated in identical ways from 100 random PSNs to obtain a fold change that was plotted. The lower and upper bound of the 95% confidence interval of the fold change was computed by dividing the actual sum of the Jaccard similarities with the 97.5th and 2.5th percentile of the random distribution, respectively, and plotted as error bars.

#### Retrieval and processing of resources for functional annotations

Co-fitness relationships between human genes were kindly provided by Joshua Pan, calculated as a Pearson correlation coefficient between the effects of the separate knockout of two genes on the AVANA cancer cell line panel^113^ as described previously^35^. Gene IDs were mapped to Ensembl gene IDs and restricted to protein-coding genes. Co-expression data on the level of correlated expression between two genes was downloaded from the SEEK^114^ database on June 13th 2018 and restricted to protein-coding genes. Pathway membership of proteins was obtained from Reactome, using a file mapping Ensembl gene IDs to lowest level pathways, downloaded on August 1st 2018. Subcellular compartment membership of proteins was obtained from the Human Protein Atlas downloaded on August 1st 2018. Annotations of ‘Uncertain’ reliability were not used. Protein complex membership of proteins was obtained from BioPlex^8^ 2.0 supplementary table 7. Entrez gene IDs were mapped to Ensembl gene IDs and restricted to protein-coding genes.

#### Calculation of significances for a network to link functionally related proteins

The significance of a given network [HuRI, HuRI 2ev (HuRI with multiple evidences), or the PSN of HuRI] to link proteins that are co-expressed or that display similar growth defects on cell lines was determined by computing for increasing correlation cutoffs the sum of the edge weights (Jaccard similarities for PSN, edge weights = 1 for PPI networks) of all edges in the given network that link two proteins with a correlation value of at least that cutoff. This sum was divided by the mean of the sums obtained from randomized networks. the 95% confidence interval was computed as described above. A single correlation cutoff (Pearson correlation coefficient of 0.3 for growth defects and co-expression value of 0.86) was chosen for display based on the number of pairs of proteins that met this cutoff. However, consistent trends were observed across most cutoffs tested (Extended Data Fig. 4d, e). The significance and confidence interval of a given network to link proteins that localize to the same subcellular compartment, or work in the same pathway, or associate in the same protein complex was computed in identical ways with the exception that no titration was performed to calculate significances.

#### Computation and visualization of functional modules

We used the SAFE software^28^ (v1.5) to determine and visualize significant functional modules in various networks. The network layouts were generated with Cytoscape^115^ (v3.4.0) using the edge-weighted spring embedded layout. PSNs were drawn using a Jaccard similarity cutoff of ≥0.1 and using the Jaccard similarity as edge weight for the layout algorithm. Gene Ontology^116^ (GO) terms for each gene were extracted from FuncAssociate^117^ (v3 – GO updated on February 2018). SAFE analysis was run with the default option except layoutAlgorithm = none (using layout generated by Cytoscape), neighborhoodRadius = 200, and neighborhoodRadiusType = absolute.

### Network coverage of genes of interest and correlation with gene properties

#### Acquisition and Processing of gene properties

The number of publications for each gene was determined as the number of unique PubMed IDs associated with the gene, using the file gene2pubmed from NCBI downloaded on August 1st 2018, after mapping NCBI gene IDs to Ensembl gene IDs, using the ID mapping file provided by NCBI, gene2ensembl, downloaded on August 1st 2018. Disease genes were defined as all genes with a disease annotation in OMIM^118^, using files generated on July 26th 2018. Pharmacological targets were taken from the IUPHAR/BPS Guide to PHARMACOLOGY^119^ downloaded on July 26th 2018. Cancer genes were the Tier 1 genes of the Cancer Gene Census^120^. Genes with Single Nucleotide Polymorphism (SNP) from Genome-Wide Association Study (GWAS) were selected using data from the GWAS Catalog^121^ v1.0.2 from July 17th 2018. Genes were selected if there was a SNP associated with a trait with *P* < 5 x 10^−8^, associated with the gene that belonged to a class that could affect the protein product. Transcription factors (TFs) were all known and likely TFs from The Human Transcription Factors^122^ v1.01 downloaded on July 26th 2018. Cell growth genes were derived from the AVANA CRISPR knockout screens^113^, using a file dated June 21st 2018 containing data for 436 cell lines. Cell growth genes were selected as those with a median relative growth < −0.5 across the cell lines tested. We took the list of genes annotated with “embryonic lethality” from the MGI database (v6.13)^34^. Transcript expression levels of genes were retrieved from GTEx, and protein expression levels from HEK293T and HeLa cell lines from these studies^9,123^. Age information was adopted from Protein Historian^36^ (http://lighthouse.ucsf.edu/ProteinHistorian, downloaded on August 2nd 2018). Age time was computed based on OrthoMCL. LoF intolerance was adopted from ExAC database^1^ (release 0.3.1 / updated February 27th 2017). Summary and detailed information of gene properties is provided in Supplementary Table 27.

#### Calculation and plotting of binned adjacency matrices

The protein-coding genome is ranked by the number of publications of each gene. In the case where multiple genes have the same number of publications, the order was randomized. The genome was divided into equal-sized bins, dividing the symmetric genome-by-genome space into two-dimensional bins.

#### Processing of gene properties for network degree correlation analysis

We calculated the correlation and the partial correlation between network degree of each network and protein properties within the set of proteins with at least one interaction in the corresponding network and with the value for the corresponding property, using Matlab (version 2016a). To calculate the partial correlation based on expression, we used HEK293T proteomic data from the BioPlex^123^ v1.0 study, proteomics data from the HeLa cell line from the QUBIC study, GTEx tissue expression data and the number of publications from NCBI (method above) for the networks BioPlex, QUBIC, CoFrac and Lit-BM, respectively.

### Integrative analyses of HuRI with subcellular compartment data

#### Calculation of protein coverage of networks by subcellular compartments

The subcellular localization dataset was retrieved from Cell Atlas^38^ and processed as described above. Any compartments with less than 100 proteins were not considered in this analysis, and additional extracellular region and extracellular vesicle annotations from the GO database were included. Fisher’s exact test was used to test for an enrichment or depletion of the proteins of each network restricted to those that have at least one localization annotation. Odds ratios were provided in log scale and colored with gray for non-significant enrichment (*P* > 0.05).

#### Calculation of trend between cellular compartment overlap and HuRI

For pairs of subcellular compartments, A and B, both the odds ratio of proteins to be annotated in both A and B and the odds ratio of the density of PPIs between proteins annotated as being in A and not B and B and not A, were calculated. The density of PPIs is the number of PPIs within a set of proteins, divided by the number of pairwise combinations of those proteins. Haldane-Anscombe corrected odds ratios were used. The uncertainties on the log odds ratios were calculated using the standard error approximation. Orthogonal distance regression (ODR) was used to estimate the relationship between the two log odds ratios. ODR was chosen since the subcellular compartments have a large variation in their sizes and so widely varying uncertainties on both the x and y variables needed to be accounted for. The z-value of the regression slope was used as the test-statistic and compared to a null distribution generated by running the same regression analysis on 1,000 degree-preserved randomized networks. All possible pairs of compartments were used, with the exclusion of pairs of compartments whose annotations partially or fully excluded each other for technical reasons. The excluded pairs were: Nucleoplasm/Nucleus, Nucleoplasm/Nucleoli, Nucleus/Nuclear Speckles, Nucleoli fibrillar center/Nucleoli, and Microtubule organizing center/Centrosome.

### Analysis of HuRI PPIs about extracellular vesicle function

#### Acquisition and processing of extracellular vesicle (EV) proteomic data from the EVpedia database

We used EVpedia database^40,124,125^ (version: April 30th 2018) to define vesicular proteins. In EVpedia, there are 487 studies for EV proteomics from human samples and an “Identification Count” for each protein, which is the number of EV proteomic studies with the corresponding protein detected. If a protein was found in more than 45 different studies (~10% of the studies), we defined it as a vesicular protein, resulting in 2,548 proteins in total (about top 10% among all the identified EV proteins). The Largest Connected Component (LCC) of this EV network, containing 525 proteins, was visualized with Cytoscape^115^ (v3.4.0). To test the significance of this EV protein network, we compared the number of PPIs between EV proteins in this network to the number of PPIs between EV proteins obtained in 1,000 degree-controlled randomized networks, deriving an empirical *P* < 0.001.

#### Design and Transfection of gRNA and selection for KO cells

We used gRNA shared in TKO database^126^. U373vIII cells were prepared at 70% confluency prior transfection. For each gene to be KO, a pool of plasmids carrying the gRNAs were transfected using JetPrime according to the manufacturer’s guidelines (Polyplus Transfection). The cells were submitted to Puromycin selection 36h after 48h of transfection. KO of SDCBP was confirmed by Western Blot^127^ with the rabbit anti-SDCBP antibody (ab133267 with lot number GR282684-7) purchased from Abcam (Cambridge).

#### Comparative proteomics

We performed the EV comparative proteomics as previously described^127^ with some modifications. The conditioned media (CM) was collected from cells grown for 72 h in culture media containing 10% EV-depleted FBS (generated by centrifugation at 150,000 g for 18 h at 4°C). CM was centrifuged at 400 g and then passed through 0.8 μm pore-size filter. The resulting filtrate was concentrated using Amicon Ultra-15 Centrifugal Filter Unit (EMD Millipore) with 100,000 nominal molecular weight limit (NMWL) molecular cutoff. The concentrate was mixed with 50% of iodixanol solution (Sigma) and processed for density gradient ultracentrifugation at 200,000 g for 2 h. The EV-enriched fraction of iodixanol was collected (at the density of ~ 1.10 g/mL) and particles were confirmed to carry CD81, an established exosome marker. The concentration of EV proteins was quantified using the BCA assay (Pierce Biotechnology). For concentration and size distribution of EVs, nanoparticle tracking analysis (NTA) was carried out with each collected iodixanol fraction using NanoSight NS500 instrument (NanoSight Ltd.). Three recordings of 30 s at 37 °C were obtained and processed using NTA software (v3.0).

The purified EV protein preparation (9 μg) was desalted with SDS-PAGE loaded onto the stacking gel followed by staining and destaining. The in-gel trypsin digestion was carried out under reducing conditions afforded by DiThioThreitol (DTT), and alkylation was achieved using iodoacetic acid. The lyophilized peptides were re-solubilized in 0.1% aqueous formic acid/2% acetonitrile, the peptides were loaded onto a Thermo Acclaim Pepmap (75 μm inner diameter with 2 cm length, C18, 3 μm particle size, 100 Å pore size; Thermo Fisher Scientific) pre-column and onto an Acclaim Pepmap Easyspray (75 μm inner diameter with 15 cm length, C18, 2 μm particle size, 100 Å pore size; Thermo Fisher Scientific) analytical column. Separation was achieved using a Dionex Ultimate 3000 uHPLC at 220 nL/min with a gradient of 2-35% organic solvents (0.1% formic acid in acetonitrile) over 3 h. Peptides were analyzed using an Orbitrap Fusion Tribrid mass spectrometer (Thermo Fisher Scientific) operating at 120,000 resolution (FWHM in MS1, 15,000 for MS/MS). All experiments were carried out in three biological replicates. The quantification of proteins was done by Maxquant program (v1.5.6.5) with default parameters of label-free quantification with UniProt database of human proteins (SwissProt release September 2016). To define the decreased proteins after knockout, *P*-value was calculated by fitting a Gaussian kernel density estimation (KDE) to the distribution of t-statistics obtained by randomly permuting the WT/KO labels for all proteins, as previously described^128^. Decreased proteins in EV after knockout were defined as the proteins with *P* < 0.05 and fold change < 0.66 (Fig. 4e). The enrichment of decreased proteins among SDCBP interactors were tested with empirical testing; among all the EV proteins from U373vIII, we randomly picked 6 proteins 10,000 times and calculated the fraction of cases with ≥ 50% decrease proteins (Extended Data Fig. 5b).

### Inference and analysis of tissue PPI networks from HuRI

#### Calculation and assessment of tissue preferential expression

A z-score related statistic published in Sonawane et al.^44^ was applied on the sample expression data of 35 tissues and testis to calculate a tissue preferential expression (TiP) value for every gene-tissue pair (Extended Data Fig. 10a and Supplementary Table 28). However, at a given TiP value cutoff, only those genes were considered preferentially expressed in a given tissue, if they also showed a level of expression > 5 in that tissue. A unique value of preferential expression for every gene was calculated by taking the maximum of all TiP values across the 35 tissues (Extended Data Fig. 10a). As noted by others^2,44^, testis by far represents the tissue with the largest number of TiP genes increasingly dominating the overall set of TiP genes for increasing TiP value cutoffs (Extended Data Fig. 10b). The expression of many genes in testis has been hypothesized to be linked to DNA integrity control mechanisms during reproduction^129^ suggesting that the majority of testis-specific genes are not related to testis-specific function. Therefore, testis was excluded from any analysis involving tissue-preferential expression in this study. The impact of that decision on the number of TiP genes defined for all other tissues was minimal, leading to a small gain in TiP genes and essentially no loss (Extended Data Fig. 10c, d). The number of TiP genes substantially decreases for increasing TiP value cutoffs (Extended Data Fig. 10a). At higher TiP value cutoffs, some tissues are left without any TiP gene. At a TiP value cutoff of 2, most TiP genes are not exclusively expressed. However, that fraction increases with higher TiP value cutoffs and plateaus at about 75% with a cutoff of 8 or 9 (Extended Data Fig. 10e). Where applicable, analyses were performed without the application of a TiP value cutoff to define TiP genes, however, whenever a cutoff was necessary, a cutoff of 2 (lenient but highly significant preferential expression), 3, and 8 were applied to test for dependencies of results on the level of preferential expression considered.

#### Correlation analysis of network properties with tissue preferential expression

Networks were processed as described above. A PPI was considered to exist in a tissue, if both proteins showed expression (at the transcript level, expression cutoff > 5) in the same sample for more than 50% of all samples available for that tissue. For the network property analyses, each PPI network was restricted to the PPIs being expressed in at least one tissue and to the largest connected component. The degree and betweenness function from the python igraph library^112^ (v0.7.1) were used to calculate the degree and betweenness of every protein in the network. Degree and betweenness were assessed for correlation with the max TiP value of every protein using Spearman’s rank correlation in the python scipy library^130^ (v1.1.0). The 95% confidence interval of Spearman’s ρ was calculated using 1,000 bootstrap samples.

#### Calculation of fraction of genome or network that is tissue-specific

Networks were processed as described above and restricted to PPIs linking proteins expressed in the same tissue. Titrations were performed as described above, by the max TiP value of every gene. The fraction of the genome that is tissue-specific at a given max TiP value cutoff was calculated by dividing the number of TiP genes at the cutoff with the number of protein-coding genes in GTEx. The fraction of proteins in a network that are tissue-specific were calculated by dividing the number of TiP proteins at a given max TiP value cutoff that are in the network with the total number of proteins in the filtered network. Error bars reflect standard error of proportion.

#### Generation and visualization of TiP networks

TiP networks were drawn for every tissue by taking all TiP proteins for a given tissue with a TiP value ≥ 2 and all their interaction partners in HuRI that are expressed in the same tissue. Networks were drawn with Cytoscape^115^ (v3.7.0) and manually adjusted for aesthetic purposes. The human body clipart was obtained from smart.servier.com.

#### Calculation of closeness of TiP proteins in tissue PPI networks

For every tissue the number of PPIs in HuRI between TiP proteins of that tissue was determined defining TiP proteins at a TiP value cutoff ≥ 2. This number was compared to the number of PPIs between TiP proteins in 1,000 degree-controlled randomized networks (generated using the degree_sequence function in the python igraph^112^ library) of the corresponding tissue PPI network of HuRI (all PPIs linking proteins expressed in this tissue). Tissues were excluded from the analysis, if less than two TiP proteins were in the tissue PPI network. The average shortest path between TiP proteins in a tissue PPI network was calculated by restricting the tissue PPI network to the Largest Connected Component (LCC), determining the shortest path of every TiP protein to the closest TiP protein and averaging those shortest paths, and comparing the average shortest path to those from 1,000 degree-controlled randomized tissue PPI networks. Tissue PPI networks with less than two TiP proteins in the LCC were excluded from the analysis. Tissues in which TiP proteins are significantly close to each other were determined by calculating the fraction of random networks with a number of TiP-TiP PPIs at least as high as in the actual tissue PPI network (or with an average shortest path at least as small or smaller) and requiring that fraction to be ≤ 0.05 × 35 (one-sided empirical p-value corrected for multiple testing).

### Gene function prediction and experimental testing

#### Analysis of annotation of TiP genes with tissue-specific GO terms

Assignment of tissues to GO^116^ terms was done manually as described in detail elsewhere (Basha et al. in preparation). These GO term – tissue annotations were further matched to the 35 tissues with transcriptome data from GTEx. GO Biological Process terms were downloaded from FuncAssociate^117^ on March 20th 2018 excluding annotations with evidence codes ND and IBA. For every tissue all genes with a TiP value ≥ 2 were selected and the fraction of those determined with at least one GO term annotation assigned to that tissue.

#### Prediction of gene functions using guilt-by-association approach

Finding a new protein annotation can be described as a link prediction problem between a node representing the function and the proteins. Initially, we connect the functional node to each of the proteins annotated with this function and obtain a link prediction score for each other gene in the network based on our recently developed link prediction method^27^. As the result, the indirect score of protein i is obtained as where aik is the connection weight between nodes i and k and kj is the degree of node j. Intuitively, the indirect score integrates the amount of network similarity of the candidate node to the known proteins involved in this function.

We then compare the original network-based indirect score (*s*) to a random benchmark, obtained by randomizing the network several times in a degree-preserved way. Calculating the *z*-score *z*=(*s-sR*)/*sR* is the traditional way of such comparison, obtained by standardizing the original score with the expectation value (*sR*) and standard deviation (*sR*) of the score that would be expected by chance. Yet, the z-score is not free from degree biases and prefers low-degree nodes with an extremely small *sR*. As described elsewhere in detail (manuscript in preparation), we propose to apply a related measure, called the effect size. The effect size *s-sR-αsR* is obtained by comparing the original score with the reasonably expected value of the random benchmark, estimated as the mean value (*sR*) and α-times the standard deviation (*sR*). In practice, we use *α*=2, selecting the same candidates as a traditional *z*-score threshold of *z* ≥ 2, but ordering them based on the amount of signal beyond random expectations to avoid a bias towards low-degree nodes. To generate the benchmark distribution for each gene, we perform 10,000 degree-preserved randomizations over the human interactome and score each gene in each of the random networks.

Functional annotations of genes with GO Biological Process terms were obtained as described above and further restricted to annotations with the evidence codes EXP, IMP, TAS, HMP, HEP, IDA, IGI, NAS, HGI, IEP, IC, HDA to avoid circularity introduced from using inferred GO annotations based on PPI data to infer more GO annotations using PPI data again.

#### Cell culture, transfection and cell death assay

Cell death assay was performed as described previously^131^. HeLa cells (obtained from American Type Culture Collection) were maintained in DMEM supplemented with 10% FBS, 0.2 mM L-glutamine, 100 U/ml penicillin and 100 μg/ml streptomycin (Invitrogen) at 37°C and 5% CO_2_. Cells were plated onto 96-well imaging plates (BD Biosciences) and transient transfection was performed using Lipofectamine 3000 according to the manufacturer’s instructions (Invitrogen). After 24h in transfection reagent, the cells were imaged on a BD Pathway 855 Bioimager (BD Biosciences) with a UAPO/340 20X objective (0.75 NA; Olympus). After 1 hour of imaging, cells were treated with 100 ng/ml of recombinant human TRAIL Apo/2L (PeproTech). The fluorescence signal and the time of death of transfected cells were calculated using ImageJ (NIH) software.

To assess the significance of a correlation between levels of OTUD6A expression and time of death, a Pearson correlation coefficient was calculated and its significance assessed by comparison to the Pearson correlation coefficients of 100,000 shufflings of the cell death and fluorescence measurements.

#### Generation of apoptosis candidate networks

Networks around OTUD6A and C6ORF222 were drawn with Cytoscape^115^ (v3.7.0). All interaction partners of OTUD6A and C6ORF222 in HuRI were selected as well as indirect interaction partners, if they had a known apoptosis GO annotation (restricted to those used in the gene function prediction, see above), to visualize the network neighborhood used to obtain the apoptosis prediction for the two candidate genes. This reference network was filtered using transcript expression data from GTEx for colon_transverse (expression cutoff > 5) and transcript expression data from the BLUEPRINT project^61,62^ for major eosinophils (log2(TPM+1) > 0.3). The same expression data was used to adjust the size of the nodes in the network to reflect transcript expression levels.

### Analyses of tissue-specificity of Mendelian diseases

#### Quantification of tissue-specificity of Mendelian diseases and integration with tissue gene expression data

Tissues affected in Mendelian diseases were manually curated as described in detail elsewhere (Basha et al. manuscript in preparation). Briefly, disease information was downloaded from the OMIM database^118^ and tissues were annotated to diseases within phenotypic series. Affected tissues were matched to the 35 tissue names from GTEx. If brain was an affected tissue, all three brain subregions for which transcriptome data from GTEx was available (see above for processing of GTEx data), were assigned to the corresponding disease. Other tissues, such as heart, were processed the same way. Annotated causal gene information from OMIM was mapped and restricted to the protein-coding gene space.

The distribution for how many diseases affect how many tissues was calculated by restricting the list of annotated Mendelian diseases and tissues to those where causal genes showed to be expressed in at least one disease-associated tissue (GTEx tissue expression > 5). Tissues were grouped into tissue types (e.g. the three brain subregions were grouped into brain) and tissue types were counted for each disease. Diseases with at most three tissue types where the disease manifests were considered tissue-specific diseases and used in all downstream analyses.

The diseases were split into those where all causal genes are TiP genes, some causal genes are TiP genes and no causal gene is a TiP gene as follows. For a given disease, every causal gene was assessed for its expression (expression > 5) in at least one of the annotated affected tissues of the given disease and whether it is preferentially expressed or not in that tissue (TiP value ≥ 2). For each disease, only causal genes that were expressed in at least one affected tissue were considered. All these causal genes for a given disease were counted based on their preferential expression and diseases grouped accordingly.

#### Test for significance of connectivity between causal proteins and TiP proteins in HuRI

To test for the significance of causal proteins of tissue-specific Mendelian diseases to interact with preferentially expressed proteins of the corresponding disease-associated diseases (hereafter referred to as disease tissues) we only considered those diseases where not a single causal proteins was found to be preferentially expressed in any of the corresponding disease tissues. Cutoffs were used as described above. For a given tissue only those HuRI PPIs were considered that linked proteins both expressed in that tissue. For every tissue, all causal genes are considered that are expressed in that tissue but that are not preferentially expressed in that tissue and that are causal to a disease that specifically manifests in that tissue and where the disease corresponds to the above mentioned criteria. No causal gene or interaction was counted twice for the analysis within each tissue. The number of PPIs between these causal proteins and TiP proteins for each tissue as well as the number of causal proteins with at least one PPI to a TiP protein was determined for each tissue and compared to the distribution of the same counts observed in 1,000 degree-controlled randomized networks that were generated with the degree_sequence function in the python igraph library.

#### Experimental design of pairwise test in Y2H to test perturbation of PPIs between causal proteins and TiP proteins by disease mutations

Causal proteins were selected as follows. Causal proteins from tissue-specific diseases were considered, if that causal protein was expressed in a given disease tissue and interacted with a TiP protein of that tissue where the causal protein itself was not preferentially expressed (cutoff as described above) and the interaction between the causal protein and TiP protein was found with the causal protein as DB fusion. For all assay versions in which a valid causal protein – TiP protein interaction was found, all interactions from these assay versions found with the corresponding causal protein as DB fusion were selected for pairwise test. Interactions involving a causal protein with 30 or more interaction partners for a given assay version were removed to control the size of the experiment. PRS v1 and RRS v1 pairs^17^ were tested along as positive and negative control.

#### Cloning of disease mutations

Within the causal genes described above, we cloned variants that were annotated as pathogenic or likely pathogenic in ClinVar^71^ (May 2015) and disease modifying in HGMD^100^ (v2016). Since then, ongoing reannotation efforts by ClinVar have classified some of these pathogenic variants as benign, conflicting or variants of uncertain significance (VUS). We generated the disease mutants by implementing an advanced high-throughput site-directed mutagenesis pipeline^68,132,133^ with some modifications as described below. For each mutation, two “primary PCRs” were performed to generate gene fragments containing the mutation and a “stitch PCR” was performed to fuse the two fragments to obtain the mutated ORF. For the primary PCRs, two universal primers (E2E forward and E2E reverse) and two ORF-specific internal forwards and reverse primers were used. The two ORF-specific primers contained the desired nucleotide change. The gene fragments generated by the primary PCRs were fused together by the stitch PCR using the universal primers to generate the mutated ORF. The final product was a full length ORF containing the mutation of interest. All the mutated ORFs were cloned into Gateway donor vector, pDONR223, by a BP reaction followed by bacterial transformation and selection by spectinomycin. Two single colonies were picked per transformant. The mutated ORF was sequence confirmed by M13 PCR followed by pooling and sequencing the PCR products on the Illumina platform. The reads were aligned to the reference ORFs using bowtie 2^103^ (v2.2.3) and samtools^134^ (v1.2). Only those colonies with the correct mutation, fully covered by the reads, and without other mutations were considered sequence-confirmed. The confirmed colonies were rearrayed and assembled followed by LR reaction and bacterial transformation to transfer the mutated ORFs in pDEST-DB. The plasmids were purified and transformed into Y8930 yeast strain for the pairwise test.

Primers used for the experiment:

E2E forward: GGCAGACGTGCCTCACTACTACAACTTTGTACAAAAAAGTTGGC

E2E reverse: CTGAGCTTGACGCATTGCTAACAACTTTGTACAAGAAAGTTGG

M13-reverse: GTAACATCAGAGATTTTGAGACAC

M13-forward: CCCAGTCACGACGTTGTAAAACG

See Supplementary Table 29 for ORF-specific primers used in the mutagenesis.

#### Pairwise test of disease mutations

All sequence-confirmed pathogenic, likely pathogenic, and reclassified alleles for all selected causal genes (see above) were subject to pairwise test along with the wild-type allele paired with all interaction partners in the respective assay versions as described above. Before the pairwise test was performed, the yeast strains containing the mutated or wild-type ORFs were Sanger sequenced to confirm the presence or absence of the mutations. Considering that all the entry clones used in the experiments have been full-length sequence verified, we were less stringent to call that a clone is sequence confirmed. Specifically, for ORFs with mutations, if either forward or reverse sanger reads confirmed the identity of the ORF and the presence of the mutations and there was no contradictory between the two reads, the clone was considered as sequence confirmed. Wild-type ORFs required one of the sanger reads confirming the identity of the ORF and absence of any of the tested mutations, and no contradictory between reads.

The pairwise test was performed in 96 well format. In all, 50 mutants in 17 genes/ORFs were subjected to pairwise test. The ORFs were inoculated in SC-Leu and SC-Trp media overnight and mated in YEPD media the following day. After incubation at 30°C overnight, the mated yeasts were transferred into SC-Leu-Trp media to select for diploids. Next day, the diploid yeasts were spotted on SC-Leu-Trp-His+3AT, SC-Leu-His+3AT+CHX and SC-Leu-Trp media to control for mating success. In parallel, we made lysates of all SC-Leu-Trp plates to perform SWIM PCR as described in the primary screening section.

#### Sequence confirmation of pairwise test positives and negatives

SWIM PCR product was sequenced and the sequencing reads were analyzed as described above in the primary screening section. Due to the short read length of the Illumina sequencing and the design of SWIM-seq, absence or presence of a mutation on the ORF sequence could not always be confirmed. Therefore, for a given pair, as long as the identities returned from the pipeline matched the tested ORF and its partner, the pair was considered as sequence confirmed. Both positives and negatives were sequenced and only combinations of sequence confirmed pairs including both wild-type and mutated ORFs with same interacting partners were included in the final analysis.

#### Processing of pairwise test data

Each spot was scored with a growth score ranging from 0 to 4, 0 being no growth, 1 being one or two colonies, 2 being some colonies, 3 being lot’s of colonies, 4 being a big fat spot where no individual colonies can be distinguished. Pairs for which the SC-Leu-Trp spot was scored as 3 or 4 and for which the CHX and the 3AT spot were valid (yeasts were spotted and no contamination or other experimental failure) were considered as successfully tested. Successfully tested pairs were further classified into auto-activators, if growth on CHX was >2 or growth on CHX was = 1 and less than 3 on 3AT, else negatives, if there was no growth (growth score = 0) on 3AT or positives if there was growth > 0 on 3AT and no growth on CHX or growth > 2 and growth = 1 on CHX. Pairs were scored blindly with respect to their identity.

#### Analysis of PPI perturbation data

Only pairs that were successfully tested, classified as positive or negative, for which the wildtype allele was classified as positive with a growth score ≥ 2, and that were sequence-confirmed were considered for all further analysis. An interaction was considered perturbed by an allele, if the growth score of the wild-type pair was at least 2 growth scores above the growth score for that interaction with the respective allele. If an interaction involving an allele was tested in multiple assay versions (because the wild-type PPI has been found in those originally), then a final decision on that interaction for being perturbed or not was based on the results from the assay version where the wild-type interaction reached the highest growth score (in case of ties, order of priority was given to assay version 1, 2, 6). A causal gene was included in the analyses, if first, there was at least one interaction with a TiP interaction partner that was positive with a growth score ≥ 2 as wild-type and that was successfully tested (as defined above) for at least one of the pathogenic alleles of that causal gene and second, if more than half of all interactions of a causal protein subjected to pairwise test were classified as positive with a growth score ≥ 2. Network visualizations were drawn with Cytoscape^115^ (v3.7.0).

### Prediction and experimental confirmation of functionally divergent splice isoforms

#### Finding tissue-regulated splicing exon

For tissue-regulated alternative splicing information, we used a dataset^78^ previously defined in 49 tissues, which was kindly shared by members of the Blencowe lab, and is now publicly available via vastDB^135^. For each of the events, we mapped Ensembl exon accession with respect to their genome coordinate information. A tissue regulated exon was defined, if there is more than 25% difference between maximum Percent Spliced-In (PSI) and minimum PSI (ΔPSI > 25) as previously described^78^.

#### Finding protein domain regulated by alternative splicing and defining hub protein with possible partial loss of protein-protein interactions

Based on the tissue-regulated exons we defined, we computationally derived the spliced-out HuRI ORFs without the corresponding exon. After that, we translated it and mapped domains for HuRI ORFs and the spliced-out HuRI ORFs with InterProScan^136^ (v5.16-55.0) with Pfam^137^ (v28.0). We restricted to domains that are previously known to mediate protein-protein interactions using information from 3did^79^ (based on UniProt version 201804). The domains that were missing in transcripts with spliced-out exons were defined as domains regulated by the alternative splicing. If a spliced-out HuRI ORF partially loses its PPI-mediating domains, we define that ORF as a possible candidate for partial PPI loss by tissue-regulated alternative splicing (Supplementary Table 18).

#### Analysis of NCK2 isoform expression in brain samples

We gained the exon-specific RPKM (Reads Per Kilobase Million) from the Allen Institute for Brain Science – BrainSpan Atlas of the Developing Human Brain^138^ (www.brainspan.org). To estimate the fraction of exon A inclusion, we divided the RPKM of exon A by the average RPKM of exon C1 and C2, which are two adjacent exons of exon A (Supplementary Table 19).

#### Pairwise test of interaction partners of NCK2 with short and long isoform

We used the orientation and Y2H assay version of the interactions where the corresponding protein interactions were found in HuRI. The pairwise test of these interactions was done as previously described^139^.

#### NCK2 function in Zebrafish

An antisense morpholino oligonucleotide, as well as a 5 nt-mismatched control morpholino oligonucleotide (Gene Tools, Inc.), was designed to knockdown expression of zNCK2B (ortholog of human NCK2) via inhibiting the removal of intron 2. Zebrafish embryos at the 1-cell stage were microinjected with 5 ng of the antisense zNCK2B or control morpholinos, and inhibition of splicing in zNCK2B was confirmed via RT-PCR at 1 dpf (days post fertilization; forward exon 2: TACGGCACAACAAGACCAGG, exon3 reverse: TTGACTATGGCCGGAGTGTT, intron2 reverse: CGTGTGCGGTCAAATTTATGC). To rescue the zNCK2B knockdown, full length and short form zNCK2B and human NCK2 (hNCK2) messenger RNA was cloned into the multiple cloning site of pCS2+MT, and transcribed with the SP6 mMessage machine kit (ThermoFisher). Purified mRNA was microinjected into 1-cell stage zebrafish embryos either alone, or in combination with the above morpholinos at a concentration sufficient to yield 0.5 fmol RNA per embryo. zNCK2B knockdown and rescued embryos were assayed at 48 hpf (hours post fertilization) for midbrain GFP expression in the zebrafish enhancer trap line SAGFF(LF)223A, where GFP was inserted adjacent to lhx9^140^. Wild-type AB embryos were similarly injected with zNCK2B or control morpholino and mRNA, fixed at 48 hpf and assayed by whole mount in situ hybridization for lef1 expression as previously described^141^. The midbrain specific lef1 expression domain was imaged and quantified by measuring the 2-dimensional area with Image-J; the experimental groups were then compared via *t*-test with MicroSoft Excel.

## Data availability

The PPI data from this publication has been submitted to the IMEx (http://www.imexconsortium.org) consortium through IntAct^106^ and assigned the identifier IM-25472. HuRI, Lit-BM, and all previously published human interactome maps from CCSB are available at http://interactome.dfci.harvard.edu/huri/ for search and download. All HuRI-related networks generated and analyzed in this study are available at NDExbio.org^142^ (https://tinyurl.com/networks-HuRI-paper). The raw and analyzed proteomic data were deposited in the PRIDE repository^143^ with the accession number PXD012321.

## Code availability

Custom code used in this study has been made available as Supplementary Data 1 for the reviewers and will be made public on github.com upon acceptance.

## References

1. Lek, M. et al. Analysis of protein-coding genetic variation in 60,706 humans. Nature 536, 285–291 (2016).

2. Melé, M. et al. The human transcriptome across tissues and individuals. Science 348, 660–665 (2015).

3. FANTOM Consortium and the RIKEN PMI and CLST (DGT). A promoter-level mammalian expression atlas. Nature 507, 462–470 (2014).

4. Regev, A. et al. The human cell atlas. eLife 6, e27041 (2017).

5. Lander, E. S. et al. Initial sequencing and analysis of the human genome. Nature 409, 860–921 (2001).

6. Venter, J. C. et al. The sequence of the human genome. Science 291, 1304–1351 (2001).

7. Wan, C. et al. Panorama of ancient metazoan macromolecular complexes. Nature 525, 339–344 (2015).

8. Huttlin, E. L. et al. Architecture of the human interactome defines protein communities and disease networks. Nature 545, 505–509 (2017).

9. Hein, M. Y. et al. A human interactome in three quantitative dimensions organized by stoichiometries and abundances. Cell 163, 712–723 (2015).

10. Rolland, T. et al. A proteome-scale map of the human interactome network. Cell 159, 1212–1226 (2014).

11. Rual, J.-F. et al. Towards a proteome-scale map of the human protein-protein interaction network. Nature 437, 1173–1178 (2005).

12. Stelzl, U. et al. A human protein-protein interaction network: a resource for annotating the proteome. Cell 122, 957–968 (2005).

13. Uhlén, M. et al. Tissue-based map of the human proteome. Science 347, 1260419 (2015).

14. Braun, P. et al. An experimentally derived confidence score for binary protein-protein interactions. Nat. Methods 6, 91–97 (2009).

15. Chen, Y.-C., Rajagopala, S. V., Stellberger, T. & Uetz, P. Exhaustive benchmarking of the yeast two-hybrid system. Nat. Methods 7, 667–668 (2010).

16. Choi, S. G. et al. Towards an “assayome” for binary interactome mapping. bioRxiv 530790 (2019).

17. Cusick, M. E. et al. Literature-curated protein interaction datasets. Nat. Methods 6, 39–46 (2009).

18. Eyckerman, S. et al. Design and application of a cytokine-receptor-based interaction trap. Nat. Cell Biol. 3, 1114–1119 (2001).

19. Cassonnet, P. et al. Benchmarking a luciferase complementation assay for detecting protein complexes. Nat. Methods 8, 990–992 (2011).

20. Berman, H. M. et al. The protein data bank. Nucleic Acids Res. 28, 235–242 (2000).

21. Mosca, R., Céol, A. & Aloy, P. Interactome3D: adding structural details to protein networks. Nat. Methods 10, 47–53 (2013).

22. Giurgiu, M. et al. CORUM: the comprehensive resource of mammalian protein complexes-2019. Nucleic Acids Res. 47, D559–D563 (2019).

23. Baccon, J., Pellizzoni, L., Rappsilber, J., Mann, M. & Dreyfuss, G. Identification and characterization of Gemin7, a novel component of the survival of motor neuron complex. J. Biol. Chem. 277, 31957–31962 (2002).

24. Tompa, P., Davey, N. E., Gibson, T. J. & Babu, M. M. A million peptide motifs for the molecular biologist. Mol. Cell 55, 161–169 (2014).

25. Leid, M. et al. Purification, cloning, and RXR identity of the HeLa cell factor with which RAR or TR heterodimerizes to bind target sequences efficiently. Cell 68, 377–395 (1992).

26. Willy, P. J. et al. LXR, a nuclear receptor that defines a distinct retinoid response pathway. Genes Dev. 9, 1033–1045 (1995).

27. Kovács, I. A. et al. Network-based prediction of protein interactions. Nat. Commun. 10, 1240 (2019).

28. Baryshnikova, A. Systematic functional annotation and visualization of biological networks. Cell Syst. 2, 412–421 (2016).

29. Graham, D. B. et al. TMEM258 is a component of the oligosaccharyltransferase complex controlling ER stress and intestinal inflammation. Cell Rep. 17, 2955–2965 (2016).

30. Yamamoto, Y., Yoshida, A., Miyazaki, N., Iwasaki, K. & Sakisaka, T. Arl6IP1 has the ability to shape the mammalian ER membrane in a reticulon-like fashion. Biochem. J. 458, 69–79 (2014).

31. Abdel-Salam, G. M. H. et al. A homozygous IER3IP1 mutation causes microcephaly with simplified gyral pattern, epilepsy, and permanent neonatal diabetes syndrome (MEDS). Am. J. Med. Genet. A. 158A, 2788–2796 (2012).

32. Yu, H. et al. High-quality binary protein interaction map of the yeast interactome network. Science 322, 104–110 (2008).

33. Jeong, H., Mason, S. P., Barabási, A. L. & Oltvai, Z. N. Lethality and centrality in protein networks. Nature 411, 41–42 (2001).

34. Smith, C. L. et al. Mouse Genome Database (MGD)-2018: knowledgebase for the laboratory mouse. Nucleic Acids Res. 46, D836–D842 (2018).

35. Pan, J. et al. Interrogation of mammalian protein complex structure, function, and membership using genome-scale fitness screens. Cell Syst. 6, 555–568.e7 (2018).

36. Capra, J. A., Williams, A. G. & Pollard, K. S. ProteinHistorian: tools for the comparative analysis of eukaryote protein origin. PLoS Comput. Biol. 8, e1002567 (2012).

37. Rito, T., Deane, C. M. & Reinert, G. The importance of age and high degree, in protein-protein interaction networks. J. Comput. Biol. 19, 785–795 (2012).

38. Thul, P. J. et al. A subcellular map of the human proteome. Science 356, (2017).

39. Youn, J.-Y. et al. High-density proximity mapping reveals the subcellular organization of mRNA-associated granules and bodies. Mol. Cell 69, 517–532 (2018).

40. Kim, D.-K. et al. EVpedia: a community web portal for extracellular vesicles research. Bioinformatics 31, 933–939 (2015).

41. Hessvik, N. P. & Llorente, A. Current knowledge on exosome biogenesis and release. Cell. Mol. Life Sci. 75, 193–208 (2018).

42. Fader, C. M., Sánchez, D. G., Mestre, M. B. & Colombo, M. I. TI-VAMP/VAMP7 and VAMP3/cellubrevin: two v-SNARE proteins involved in specific steps of the autophagy/multivesicular body pathways. Biochim. Biophys. Acta 1793, 1901–1916 (2009).

43. Imjeti, N. S. et al. Syntenin mediates SRC function in exosomal cell-to-cell communication. Proc. Natl. Acad. Sci. U. S. A. 114, 12495–12500 (2017).

44. Sonawane, A. R. et al. Understanding tissue-specific gene regulation. Cell Rep. 21, 1077–1088 (2017).

45. Calderone, A., Castagnoli, L. & Cesareni, G. Mentha: a resource for browsing integrated protein-interaction networks. Nat. Methods 10, 690–691 (2013).

46. Barabási, A.-L. & Oltvai, Z. N. Network biology: understanding the cell’s functional organization. Nat. Rev. Genet. 5, 101–113 (2004).

47. Kiran, M. & Nagarajaram, H. A. Global versus local hubs in human protein-protein interaction network. J. Proteome Res. 12, 5436–5446 (2013).

48. Lin, W.-H., Liu, W.-C. & Hwang, M.-J. Topological and organizational properties of the products of house-keeping and tissue-specific genes in protein-protein interaction networks. BMC Syst. Biol. 3, 32 (2009).

49. Yang, L. et al. Comparative analysis of housekeeping and tissue-selective genes in human based on network topologies and biological properties. Mol. Genet. Genomics 291, 1227–1241 (2016).

50. Paulson, J. N. et al. Tissue-aware RNA-Seq processing and normalization for heterogeneous and sparse data. BMC Bioinformatics 18, 437 (2017).

51. Bossi, A. & Lehner, B. Tissue specificity and the human protein interaction network. Mol. Syst. Biol. 5, 260 (2009).

52. Singh, R., Letai, A. & Sarosiek, K. Regulation of apoptosis in health and disease: the balancing act of BCL-2 family proteins. Nat. Rev. Mol. Cell Biol. 20, 175–193 (2019).

53. Bouillet, P. et al. Proapoptotic Bcl-2 relative Bim required for certain apoptotic responses, leukocyte homeostasis, and to preclude autoimmunity. Science 286, 1735–1738 (1999).

54. Huang, S., Tang, R. & Poon, R. Y. C. BCL-W is a regulator of microtubule inhibitor-induced mitotic cell death. Oncotarget 7, 38718–38730 (2016).

55. Chao, D. T. et al. Bcl-XL and Bcl-2 repress a common pathway of cell death. J. Exp. Med. 182, 821–828 (1995).

56. Xu, G. et al. Lipocalin-2 induces cardiomyocyte apoptosis by increasing intracellular iron accumulation. J. Biol. Chem. 287, 4808–4817 (2012).

57. D’Sa-Eipper, C. & Chinnadurai, G. Functional dissection of Bfl-1, a Bcl-2 homolog: antiapoptosis, oncogene-cooperation and cell proliferation activities. Oncogene 16, 3105–3114 (1998).

58. Zhai, D. et al. Characterization of the anti-apoptotic mechanism of Bcl-B. Biochem. J. 376, 229–236 (2003).

59. Puthalakath, H. et al. Bmf: a proapoptotic BH3-only protein regulated by interaction with the myosin V actin motor complex, activated by anoikis. Science 293, 1829–1832 (2001).

60. Puthalakath, H., Huang, D. C., O’Reilly, L. A., King, S. M. & Strasser, A. The proapoptotic activity of the Bcl-2 family member Bim is regulated by interaction with the dynein motor complex. Mol. Cell 3, 287–296 (1999).

61. Adams, D. et al. BLUEPRINT to decode the epigenetic signature written in blood. Nat. Biotechnol. 30, 224–226 (2012).

62. Stunnenberg, H. G., International Human Epigenome Consortium & Hirst, M. The International Human Epigenome Consortium: a blueprint for scientific collaboration and discovery. Cell 167, 1145–1149 (2016).

63. Petryszak, R. et al. Expression Atlas update - an integrated database of gene and protein expression in humans, animals and plants. Nucleic Acids Res. 44, D746–752 (2016).

64. Purcell, M., Kruger, A. & Tainsky, M. A. Gene expression profiling of replicative and induced senescence. Cell Cycle 13, 3927–3937 (2014).

65. DeBartolo, J., Taipale, M. & Keating, A. E. Genome-wide prediction and validation of peptides that bind human prosurvival Bcl-2 proteins. PLoS Comput. Biol. 10, e1003693 (2014).

66. Barshir, R. et al. Role of duplicate genes in determining the tissue-selectivity of hereditary diseases. PLoS Genet. 14, e1007327 (2018).

67. Barshir, R., Shwartz, O., Smoly, I. Y. & Yeger-Lotem, E. Comparative analysis of human tissue interactomes reveals factors leading to tissue-specific manifestation of hereditary diseases. PLoS Comput. Biol. 10, e1003632 (2014).

68. Sahni, N. et al. Widespread macromolecular interaction perturbations in human genetic disorders. Cell 161, 647–660 (2015).

69. Shen, J. et al. Mutations in PNKP cause microcephaly, seizures and defects in DNA repair. Nat. Genet. 42, 245–249 (2010).

70. Reynolds, J. J., Walker, A. K., Gilmore, E. C., Walsh, C. A. & Caldecott, K. W. Impact of PNKP mutations associated with microcephaly, seizures and developmental delay on enzyme activity and DNA strand break repair. Nucleic Acids Res. 40, 6608–6619 (2012).

71. Landrum, M. J. et al. ClinVar: improving access to variant interpretations and supporting evidence. Nucleic Acids Res. 46, D1062–D1067 (2018).

72. Bhatnagar, S. et al. TRIM37 is a new histone H2A ubiquitin ligase and breast cancer oncoprotein. Nature 516, 116–120 (2014).

73. Olivé, M. et al. New cardiac and skeletal protein aggregate myopathy associated with combined MuRF1 and MuRF3 mutations. Hum. Mol. Genet. 24, 3638–3650 (2015).

74. Novarino, G. et al. Exome sequencing links corticospinal motor neuron disease to common neurodegenerative disorders. Science 343, 506–511 (2014).

75. Corominas, R. et al. Protein interaction network of alternatively spliced isoforms from brain links genetic risk factors for autism. Nat. Commun. 5, 3650 (2014).

76. Yang, X. et al. Widespread expansion of protein interaction capabilities by alternative splicing. Cell 164, 805–817 (2016).

77. Yeo, G., Holste, D., Kreiman, G. & Burge, C. B. Variation in alternative splicing across human tissues. Genome Biol. 5, R74 (2004).

78. Irimia, M. et al. A highly conserved program of neuronal microexons is misregulated in autistic brains. Cell 159, 1511–1523 (2014).

79. Mosca, R., Céol, A., Stein, A., Olivella, R. & Aloy, P. 3did: a catalog of domain-based interactions of known three-dimensional structure. Nucleic Acids Res. 42, D374–379 (2014).

80. Bladt, F. et al. The murine Nck SH2/SH3 adaptors are important for the development of mesoderm-derived embryonic structures and for regulating the cellular actin network. Mol. Cell. Biol. 23, 4586–4597 (2003).

81. Ngoenkam, J. et al. Non-overlapping functions of Nck1 and Nck2 adaptor proteins in T cell activation. Cell Commun. Signal. 12, 21 (2014).

82. Thévenot, E. et al. P21-Activated Kinase 3 (PAK3) protein regulates synaptic transmission through its interaction with the Nck2/Grb4 protein adaptor. J. Biol. Chem. 286, 40044–40059 (2011).

83. Fawcett, J. P. et al. Nck adaptor proteins control the organization of neuronal circuits important for walking. Proc. Natl. Acad. Sci. U. S. A. 104, 20973–20978 (2007).

84. Wegmeyer, H. et al. EphA4-dependent axon guidance is mediated by the RacGAP alpha2-chimaerin. Neuron 55, 756–767 (2007).

85. Cowan, C. A. & Henkemeyer, M. The SH2/SH3 adaptor Grb4 transduces B-ephrin reverse signals. Nature 413, 174–179 (2001).

86. Hyung, D., Kim, J., Cho, S. Y. & Park, C. ASpedia: a comprehensive encyclopedia of human alternative splicing. Nucleic Acids Res. 46, D58–D63 (2018).

87. Xiong, H. Y. et al. RNA splicing. The human splicing code reveals new insights into the genetic determinants of disease. Science 347, 1254806 (2015).

88. Celaj, A. et al. Quantitative analysis of protein interaction network dynamics in yeast. Mol. Syst. Biol. 13, 934 (2017).

89. Skinnider, M. A. et al. An atlas of protein-protein interactions across mammalian tissues. bioRxiv 351247 (2018).

90. Yachie, N. et al. Pooled-matrix protein interaction screens using Barcode Fusion Genetics. Mol. Syst. Biol. 12, 863 (2016).

91. Weile, J. et al. A framework for exhaustively mapping functional missense variants. Mol. Syst. Biol. 13, 957 (2017).

## References

92. Yang, X. et al. A public genome-scale lentiviral expression library of human ORFs. Nat. Methods 8, 659–661 (2011).

93. ORFeome Collaboration. The ORFeome Collaboration: a genome-scale human ORF-clone resource. Nat. Methods 13, 191–192 (2016).

94. Seiler, C. Y. et al. DNASU plasmid and PSI:biology-materials repositories: resources to accelerate biological research. Nucleic Acids Res. 42, D1253–1260 (2014).

95. Kent, W. J. BLAT – the BLAST-like alignment tool. Genome Res. 12, 656–664 (2002).

96. O’Leary, N. A. et al. Reference sequence (RefSeq) database at NCBI: current status, taxonomic expansion, and functional annotation. Nucleic Acids Res. 44, D733–745 (2016).

97. Frankish, A. et al. GENCODE reference annotation for the human and mouse genomes. Nucleic Acids Res. 47, D766–D773 (2019).

98. Altschul, S. F., Gish, W., Miller, W., Myers, E. W. & Lipman, D. J. Basic local alignment search tool. J. Mol. Biol. 215, 403–410 (1990).

99. Edgar, R. C. MUSCLE: multiple sequence alignment with high accuracy and high throughput. Nucleic Acids Res. 32, 1792–1797 (2004).

100. Stenson, P. D. et al. Human Gene Mutation Database (HGMD): 2003 update. Hum. Mutat. 21, 577–581 (2003).

101. Stellberger, T. et al. Improving the yeast two-hybrid system with permutated fusions proteins: the Varicella Zoster Virus interactome. Proteome Sci. 8, 8 (2010).

102. Dreze, M. et al. High-quality binary interactome mapping. Methods Enzymol. 470, 281–315 (2010).

103. Langmead, B. & Salzberg, S. L. Fast gapped-read alignment with Bowtie 2. Nat. Methods 9, 357–359 (2012).

104. Lawless, C., Wilkinson, D. J., Young, A., Addinall, S. G. & Lydall, D. A. Colonyzer: automated quantification of micro-organism growth characteristics on solid agar. BMC Bioinformatics 11, 287 (2010).

105. Licata, L. et al. MINT, the molecular interaction database: 2012 update. Nucleic Acids Res. 40, D857–861 (2012).

106. Orchard, S. et al. The MlntAct project – IntAct as a common curation platform for 11 molecular interaction databases. Nucleic Acids Res. 42, D358–363 (2014).

107. Salwinski, L. et al. The database of interacting proteins: 2004 update. Nucleic Acids Res. 32, D449–451 (2004).

108. Launay, G., Salza, R., Multedo, D., Thierry-Mieg, N. & Ricard-Blum, S. MatrixDB, the extracellular matrix interaction database: updated content, a new navigator and expanded functionalities. Nucleic Acids Res. 43, D321–327 (2015).

109. Chatr-Aryamontri, A. et al. The BioGRID interaction database: 2017 update. Nucleic Acids Res. 45, D369–D379 (2017).

110. Kerrien, S. et al. Broadening the horizon – level 2.5 of the HUPO-PSI format for molecular interactions. BMC Biol. 5, 44 (2007).

111. Ono, K., Muetze, T., Kolishovski, G., Shannon, P. & Demchak, B. CyREST: turbocharging Cytoscape access for external tools via a RESTful API. F1000Research 4, 478 (2015).

112. Csardi, G. & Nepusz, T. The igraph software package for complex network research. InterJournal Complex Syst. 1695, 1–9 (2006).

113. Tsherniak, A. et al. Defining a cancer dependency map. Cell 170, 564–576 (2017).

114. Zhu, Q. et al. Targeted exploration and analysis of large cross-platform human transcriptomic compendia. Nat. Methods 12, 211–214 (2015).

115. Shannon, P. et al. Cytoscape: a software environment for integrated models of biomolecular interaction networks. Genome Res. 13, 2498–2504 (2003).

116. Ashburner, M. et al. Gene ontology: tool for the unification of biology. Nat. Genet. 25, 25–29 (2000).

117. Berriz, G. F., King, O. D., Bryant, B., Sander, C. & Roth, F. P. Characterizing gene sets with FuncAssociate. Bioinformatics 19, 2502–2504 (2003).

118. Amberger, J. S., Bocchini, C. A., Schiettecatte, F., Scott, A. F. & Hamosh, A. OMIM.org: Online Mendelian Inheritance in Man (OMIM®), an online catalog of human genes and genetic disorders. Nucleic Acids Res. 43, D789–798 (2015).

119. Harding, S. D. et al. The lUPHAR/BPS Guide to PHARMACOLOGY in 2018: updates and expansion to encompass the new guide to IMMUNOPHARMACOLOGY. Nucleic Acids Res. 46, D1091–D1106 (2018).

120. Tate, J. G. et al. COSMIC: the Catalogue Of Somatic Mutations In Cancer. Nucleic Acids Res. 47, D941–D947 (2019).

121. Buniello, A. et al. The NHGRI-EBI GWAS Catalog of published genome-wide association studies, targeted arrays and summary statistics 2019. Nucleic Acids Res. 47, D1005–D1012 (2019).

122. Lambert, S. A. et al. The human transcription factors. Cell 172, 650–665 (2018).

123. Huttlin, E. L. et al. The BioPlex network: a systematic exploration of the human interactome. Cell 162, 425–440 (2015).

124. Kim, D.-K. et al. EVpedia: an integrated database of high-throughput data for systemic analyses of extracellular vesicles. J. Extracell. Vesicles 2, (2013).

125. Kim, D.-K., Lee, J., Simpson, R. J., Lötvall, J. & Gho, Y. S. EVpedia: A community web resource for prokaryotic and eukaryotic extracellular vesicles research. Semin. Cell Dev. Biol. 40, 4–7 (2015).

126. Hart, T. et al. Evaluation and design of genome-wide CRISPR/SpCas9 knockout screens. G3 7, 2719–2727 (2017).

127. Choi, D. et al. The impact of oncogenic EGFRvlll on the proteome of extracellular vesicles released from glioblastoma cells. Mol. Cell. Proteomics 17, 1948–1964 (2018).

128. Kim, H. J. et al. Time-evolving genetic networks reveal a NAC troika that negatively regulates leaf senescence in Arabidopsis. Proc. Natl. Acad. Sci. U. S. A. 115, E4930–E4939 (2018).

129. Xia, B. et al. Widespread transcriptional scanning in the testis modulates gene evolution rates. bioRxiv 282129 (2019).

130. Jones, E., Oliphant, T. & Peterson, P. SciPy: open source scientific tools for python. (2001). Available at: http://www.scipy.org/. (Accessed: 3rd April 2019)

131. Lee, R. E. C., Walker, S. R., Savery, K., Frank, D. A. & Gaudet, S. Fold-change of nuclear NF-kB determines TNF-induced transcription in single cells. Mol. Cell 53, 867–879 (2014).

132. Zhong, Q. et al. Edgetic perturbation models of human inherited disorders. Mol. Syst. Biol. 5, 321 (2009).

133. Charloteaux, B. et al. Protein-protein interactions and networks: forward and reverse edgetics. Methods Mol. Biol. 759, 197–213 (2011).

134. Li, H. et al. The Sequence Alignment/Map format and SAMtools. Bioinformatics 25, 2078–2079 (2009).

135. Tapial, J. et al. An atlas of alternative splicing profiles and functional associations reveals new regulatory programs and genes that simultaneously express multiple major isoforms. Genome Res. 27, 1759–1768 (2017).

136. Jones, P. et al. InterProScan 5: genome-scale protein function classification. Bioinformatics 30, 1236–1240 (2014).

137. El-Gebali, S. et al. The Pfam protein families database in 2019. Nucleic Acids Res. 47, D427–D432 (2019).

138. Miller, J. A. et al. Transcriptional landscape of the prenatal human brain. Nature 508, 199–206 (2014).

139. Betts, M. J. et al. Systematic identification of phosphorylation-mediated protein interaction switches. PLoS Comput. Biol. 13, e1005462 (2017).

140. Kawakami, K. et al. zTrap: zebrafish gene trap and enhancer trap database. BMC Dev. Biol. 10, 105 (2010).

141. Olsen, J. B. et al. G9a and ZNF644 physically associate to suppress progenitor gene expression during neurogenesis. Stem Cell Rep. 7, 454–470 (2016).

142. Pratt, D. et al. NDEx, the Network Data Exchange. Cell Syst. 1, 302–305 (2015).

143. Perez-Riverol, Y. et al. The PRIDE database and related tools and resources in 2019: improving support for quantification data. Nucleic Acids Res. 47, D442–D450 (2019).

