## Extended Data Figures for "A reference map of the human protein interactome"

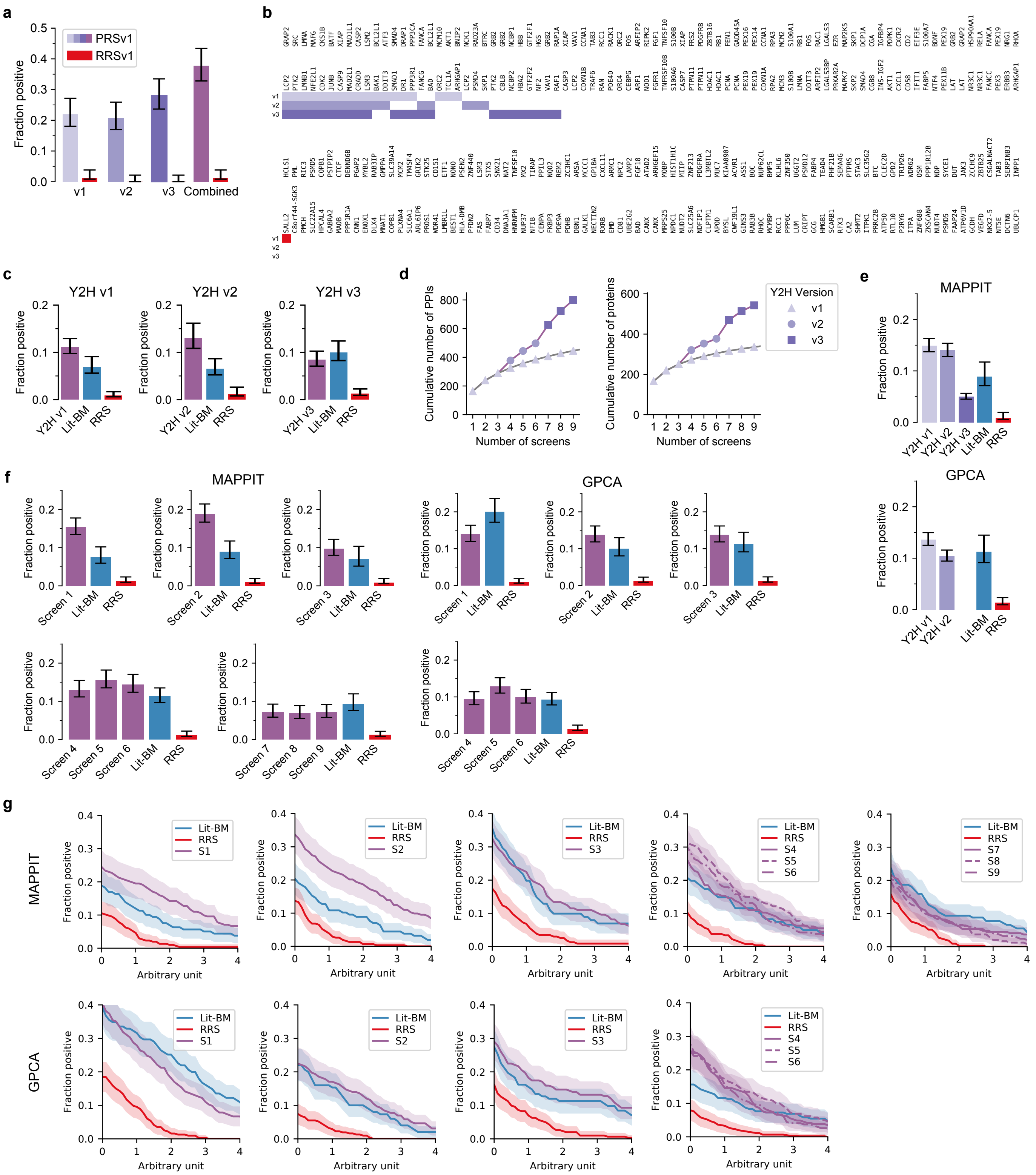

**Extended Data Fig. 1 | Y2H assay development and validation of HuRI.** **a,** Individual and combined recovery of PRSv1 and RRSv1 pairs by Y2H assay versions. **b,** Colored squares showing which protein pairs were detected in PRSv1 (top) and RRSv1 (bottom) by Y2H assay versions. **c,** Recovery rates of Lit-BM and PPIs from screens of a 2k-by-2k gene test space per Y2H assay version in MAPPIT. **d,** Cumulative PPI and protein count performing three screens with each Y2H assay version in the test space compared to nine screens with Y2H assay version 1. **e,f,** Recovery of Lit-BM and PPIs from screens of Space III when split by assay version (**e**) or screen at an RRS rate of 1% (**f**) or across a range of thresholds (**g**) in MAPPIT and GPCA. All error bars, in **a**, **c**, **e**, **f**, are 68.3% Bayesian CI.

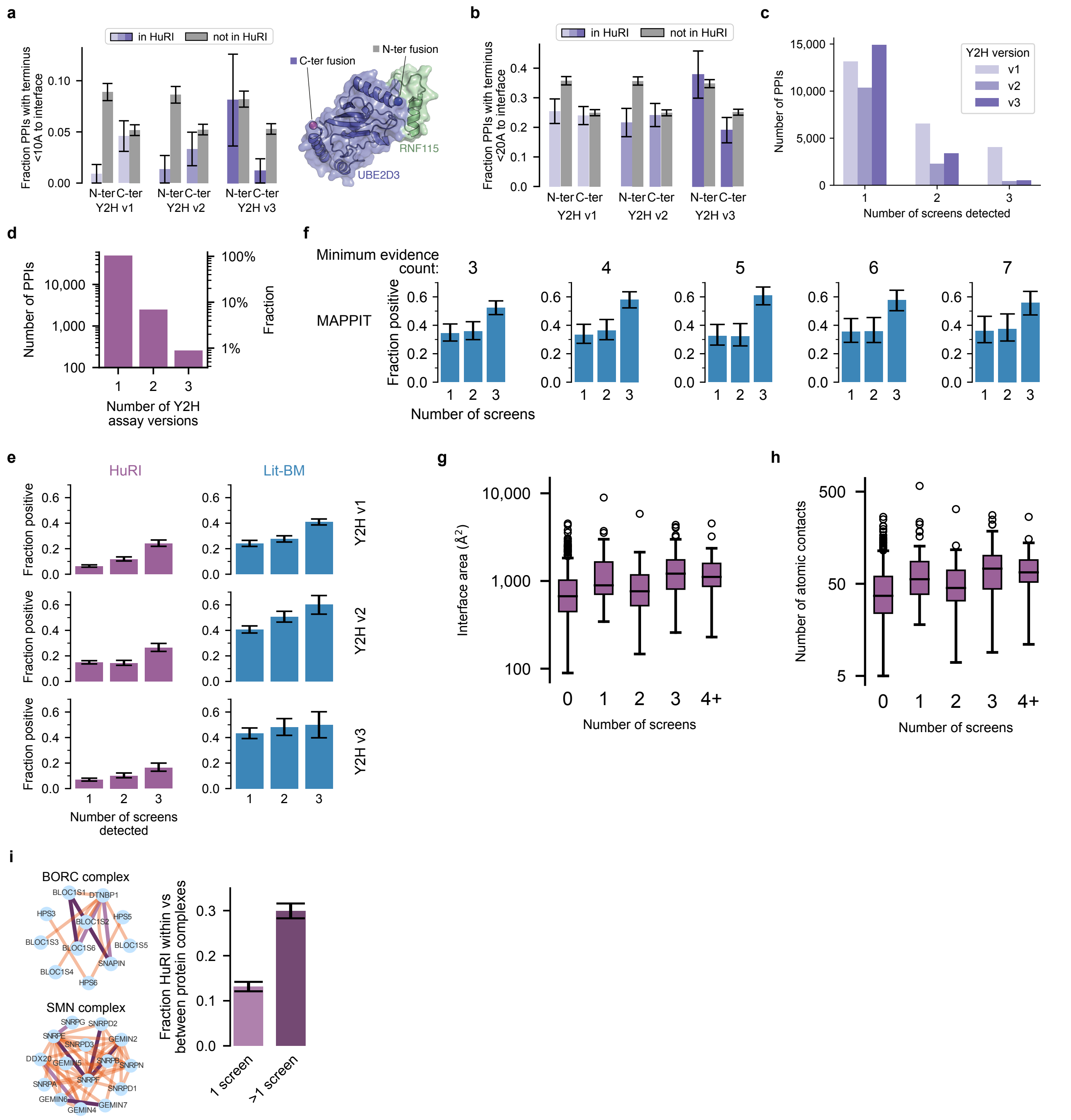


**Extended Data Fig. 2 | Stericity and interaction strength contribute to PPI detectability. a, b,** Fraction of PPIs with N or C-terminus < 10 Å (**a**) or 20 Å (**b**) to PPI interface, for PPIs with known structure in and not in HuRI. Error bars are standard error of proportion. The structure of UBE2D3 bound to RNF115 illustrates an example of a PPI found only by Y2H assay version 3 (PDB code: 5ulh). **c** Number of screens each PPI in HuRI was detected in, split by Y2H assay version. **d,** Number of Y2H assay versions each PPI in HuRI was detected in. **e,** MAPPIT recovery rates of HuRI and Lit-BM PPIs that were also detected in HuRI by the number of screens each pair was detected in. Error bars are 68.3% Bayesian CI. **f,** MAPPIT recovery rates of Lit-BM PPIs that were also detected in HuRI, for increasing number of pieces of experimental evidence per PPI. Error bars are 68.3% Bayesian CI. **g/h,** The size of the interaction interface area (**g**) / number of atomic contacts (**h**) by the number of HuRI screens in which a PPI is detected, with boxplots showing median, interquartile range (IQR), and 1.5 x IQR (with outliers). **i,** Examples of within-complex interactions detected in HuRI (purple) and BioPlex (orange). Fraction of HuRI PPIs between proteins of protein complexes that link proteins of the same complex, split by PPIs found in single and multiple screens (dark purple). Error bars are standard error of proportion.

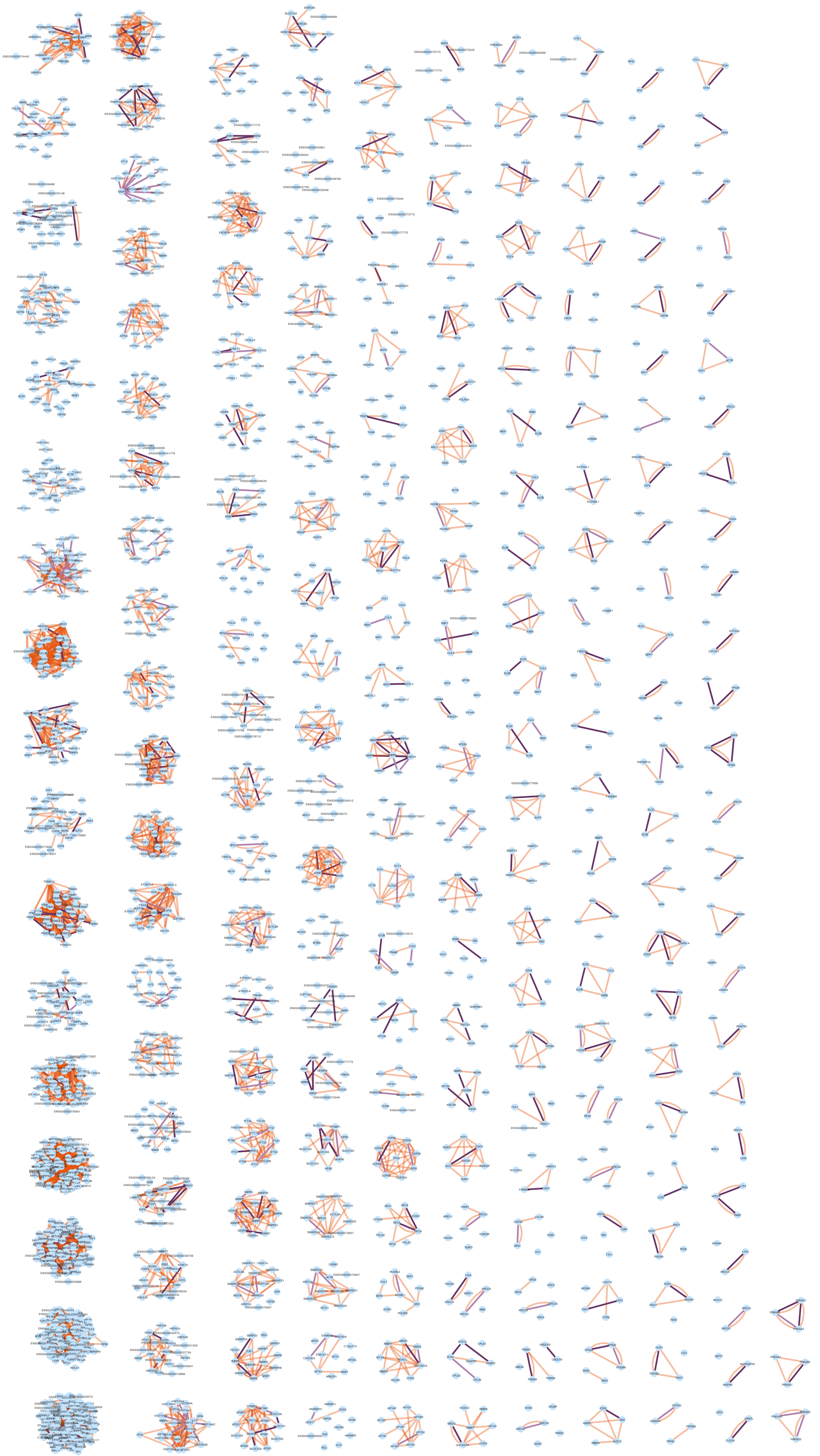

**Extended Data Fig. 3 | HuRI provides direct contact information for proteins in complexes.** Intra-complex PPIs are shown for protein complexes from CORUM as found in BioPlex (orange) or HuRI (purple). HuRI PPIs are further distinguished into PPIs found in single and multiple screens (dark purple).

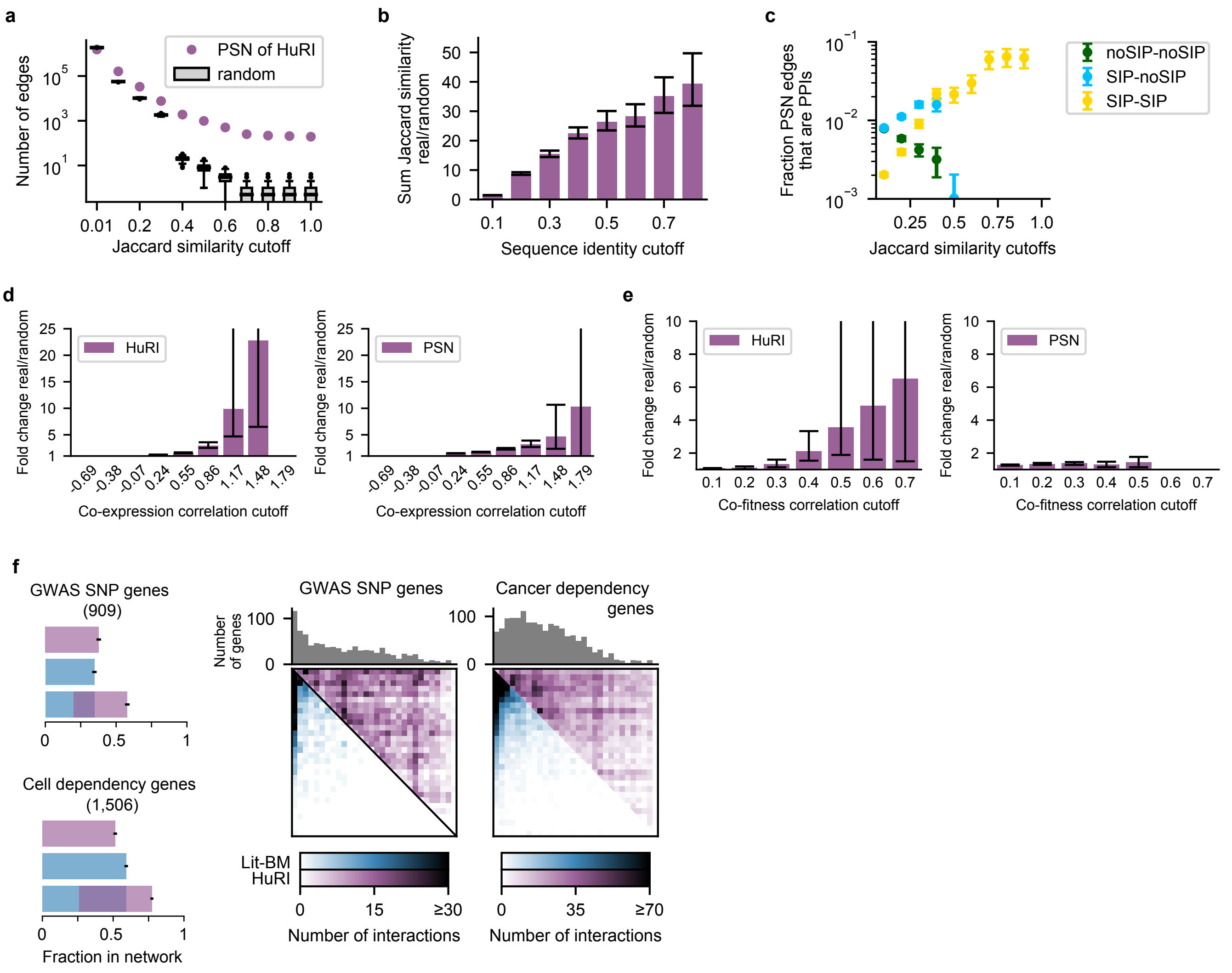


**Extended Data Fig. 4 | Topological and functional significance of HuRI.** **a,** The number of pairs of proteins in HuRI and random networks at increasing Jaccard similarity cutoffs, with boxplots showing median, interquartile range (IQR), and 1.5 x IQR (with outliers). PSN: profile similarity network. **b,** Fold change over random networks of the sum of Jaccard similarities of pairs of proteins in HuRI above at increasing thresholds of sequence identity. Error bars are 95% confidence intervals. **c,** Fraction of PSN edges that are also PPIs in HuRI, split by the PPIs involving no, one or two self-interacting proteins (SIPs), at increasing Jaccard similarity cutoffs. Error bars are standard error of proportion. **d**, **e**, Fold change over random networks of the PPI count (left) or sum of Jaccard similarities (right) of HuRI PPIs or PSN pairs, respectively, at increasing co-expression (**d**) and co-fitness (**e**) cutoffs. Error bars are 95% confidence interval. **f**, Fraction of genes with at least one PPI for biomedically interesting genes. Heatmap of HuRI and Lit-BM PPI counts between proteins, ordered by number of publications, restricted to PPIs involving genes from the corresponding gene set.

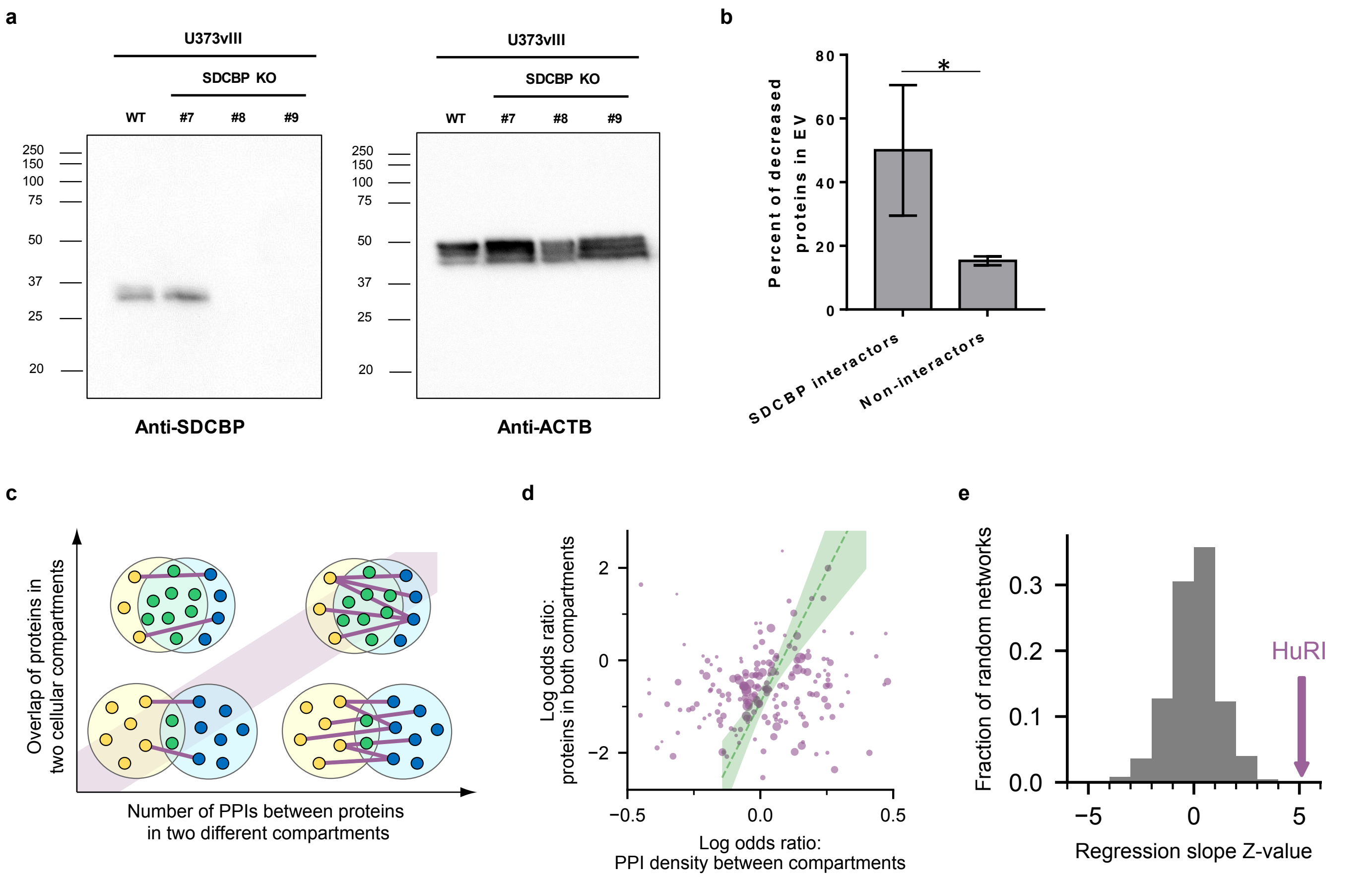


**Extended Data Fig. 5 | Incomplete protein localization annotation likely underlies apparent lack of co-localization of proteins interacting in HuRI.** **a,** Western Blot of SDCBP (left panel) and ACTB (loading control, right panel) in wild-type (WT) and three knockout (KO) cell lines (#7-#9). Cell line #8 was used for EV proteomics. **b**, Fraction of proteins whose abundance in EVs was significantly reduced in the SDCBP KO cell line, split by proteins interacting and not interacting with SDCBP as identified in HuRI. Error bars are standard error of proportion. **c,** Schematic illustrating that the number of HuRI PPIs between proteins from two different compartments should correlate with the enrichment of both compartment pairs to overlap, if co-localization annotation is incomplete. **d,** Scatter plot showing, for each pair of subcellular compartments, odds ratios quantifying the enrichment for proteins located in both compartments versus the enrichment of the density of PPIs between proteins located to either compartment. Size of points is scaled by the standard error of the x axis variable. Regression line and 95% confidence interval are shown. **e,** The z-score of the regression slope of **g** compared to those of random networks.

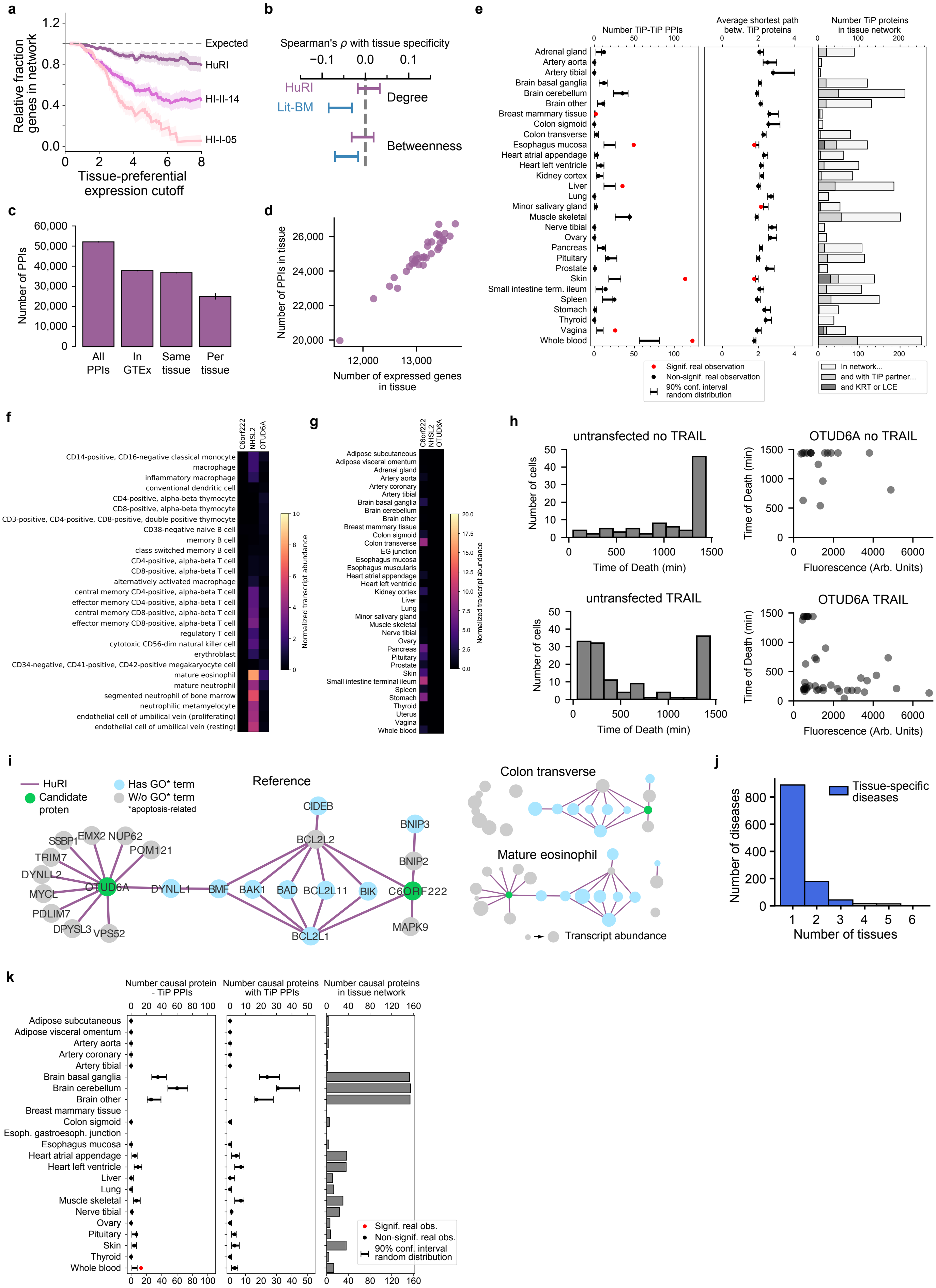
a
Relative fraction genes in network
Expected
HuRI
HI-II-14
HI-I-05
Tissue-preferential expression cutoff
b
Spearman's ρ with tissue specificity
HuRI
Lit-BM
Degree
Betweenness
c
Number of PPIs
All PPIs
In GTEx
Same tissue
Per tissue
d
Number of PPIs in tissue
Number of expressed genes in tissue
e
Number TiP-TiP PPIs
Average shortest path betw. TiP proteins
Number TiP proteins in tissue network
Adrenal gland
Artery aorta
Artery tibial
Brain basal ganglia
Brain cerebellum
Brain other
Breast mammary tissue
Colon sigmoid
Colon transverse
Esophagus mucosa
Heart atrial appendage
Heart left ventricle
Kidney cortex
Liver
Lung
Minor salivary gland
Muscle skeletal
Nerve tibial
Ovary
Pancreas
Pituitary
Prostate
Skin
Small intestine term. ileum
Spleen
Stomach
Thyroid
Vagina
Whole blood
Signif. real observation
Non-signif. real observation
90% conf. interval random distribution
In network...
and with TiP partner...
and KRT or LCE
f
C6orf222
NHSL2
OTUD6A
Normalized transcript abundance
g
Normalized transcript abundance
h
untransfected no TRAIL
OTUD6A no TRAIL
untransfected TRAIL
OTUD6A TRAIL
Number of cells
Time of Death (min)
Fluorescence (Arb. Units)
i
HuRI
Candidate proten
Has GO* term
W/o GO* term
*apoptosis-related
Reference
Colon transverse
Mature eosinophil
Transcript abundance
j
Number of diseases
Tissue-specific diseases
Number of tissues
k
Number causal protein - TiP PPIs
Number causal proteins with TiP PPIs
Number causal proteins in tissue network
Signif. real obs.
Non-signif. real obs.
90% conf. interval random distribution

**Extended Data Fig. 6 | PPIs between TiP proteins and uniformly expressed proteins likely adapt basic cellular processes to mediate cellular context-specific functions.** **a**, TiP protein coverage by CCSB PPI networks for increasing levels of tissue-preferential expression. **b,** Spearman correlation coefficients and 95% confidence intervals for correlations between degree or betweenness and tissue specificity for HuRI and Lit-BM. **c**, Number of PPIs in HuRI, involving proteins in GTEx, where both proteins are expressed in the same tissue, and the mean of the tissue-specific subnetworks where error bar is standard deviation. **d,** Scatterplot showing the number of HuRI PPIs in each tissue network as a function of the number of genes expressed in the tissue. **e,** Test for enrichment of TiP-TiP PPIs (left) and significance of average shortest path between TiP proteins (middle) in each tissue subnetwork, number of TiP proteins in each subnetwork, interacting with other TiP proteins, being part of Keratin (KRT) or Late-cornified envelope (LCE) protein family (right). **f, g,** Transcript expression levels across the GTEx tissue panel (**f**) and BLUEPRINT hematopoietic cell lineage (**g**) for three candidate genes predicted to function in apoptosis. EG = esophagus gastroesophageal. **h,** Histogram of number of untransfected cells and their time of death without and with addition of TRAIL (left). Time of death of cells expressing OTUD6A-GFP fusions versus OTUD6A expression measured as fluorescence without and with addition of TRAIL (right). **i,** Apoptosis-related network context of OTUD6A and C6ORF222 in HuRI, unfiltered (left) and filtered using colon transverse or mature eosinophil transcript levels (right). **j,** Histogram of the number of Mendelian diseases showing symptoms in a number of tissues. **k,** Test for enrichment of causal proteins associated with tissue-specific Mendelian diseases to interact with TiP proteins of affected tissues.

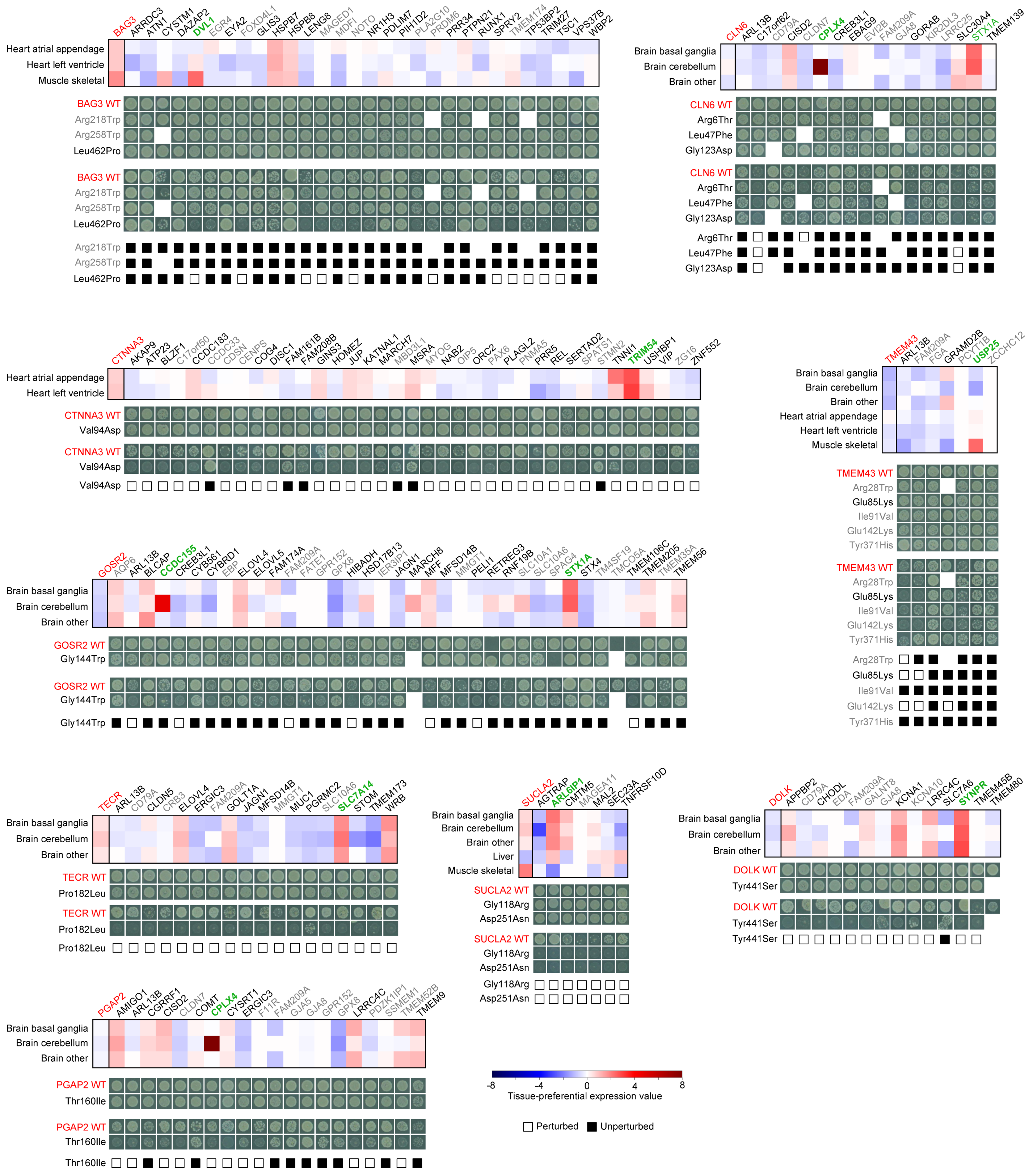


**Extended Data Fig. 7 | Mutations in uniformly expressed causal proteins associated with tissue-specific Mendelian diseases perturb interactions to TiP proteins.** Expression profile and interaction perturbation profile of nine causal proteins and their interaction partners. Affected tissues were selected for display (top). Control of AD and DB plasmid presence and cell density by spotting yeast colonies on SC-Leu-TRP media (upper). Detection of PPIs by spotting yeast on SC-Leu-Trp-His+3AT media (lower). Yeast growth indicates PPIs. WT = wild-type, red letters = causal proteins or alleles, grey gene symbols = interaction partners not expressed in affected tissues, grey alleles = not pathogenic, green gene symbols = TiP interaction partners in affected tissues.

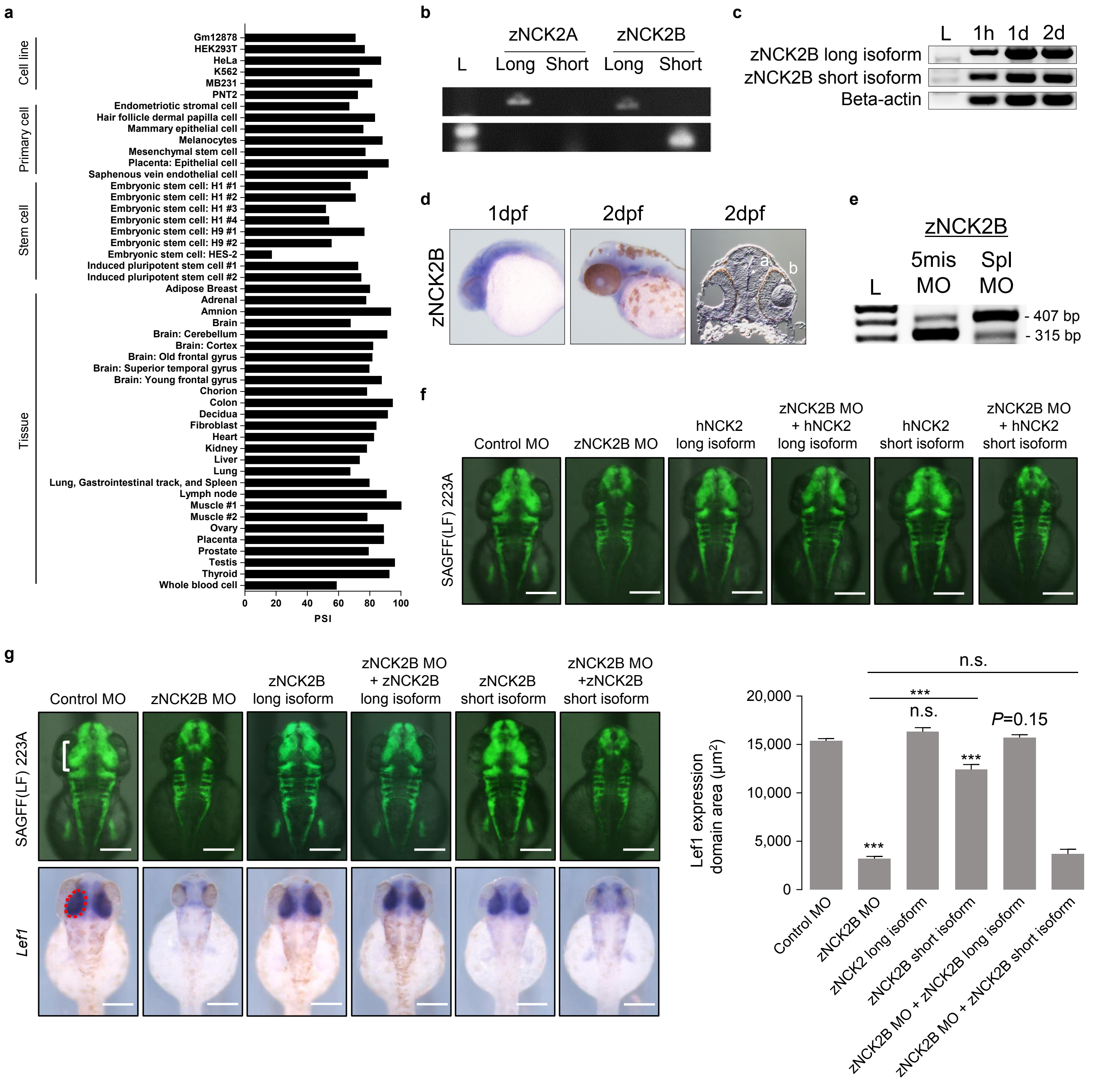


**Extended Data Fig. 8 | Experimental characterization of the dominant negative effect of the short isoform of NCK2 in a zebrafish model.** **a,** Fraction of NCK2 transcripts with alternative exon spliced in (long isoform) across tissues and cell types. PSI: percent spliced in. **b,** RT-PCR analysis of isoforms of the two NCK2 orthologs in zebrafish. **c,** RT-PCR analysis on transcript expression of zNCK2B at various time points. L: size marker lane; 1h: 1 hour after fertilization (maternal expression); 1d, 2d: 1 and 2 days of development. **d,** Whole mount *in situ* hybridization of zNCK2B in the anterior central nervous system, including domains in the midbrain and retina (white arrowheads). dpf = days post-fertilization. **e,** Splicing regulation by antisense splice MO (5ng) but not by 5-mismatch control MO. L = size marker lane. **f,** *In vivo* test of distinct functions of human NCK2 isoforms with different marker. **g,** *In vivo* test of distinct functions of zebrafish NCK2B isoforms. Error bars are standard error.

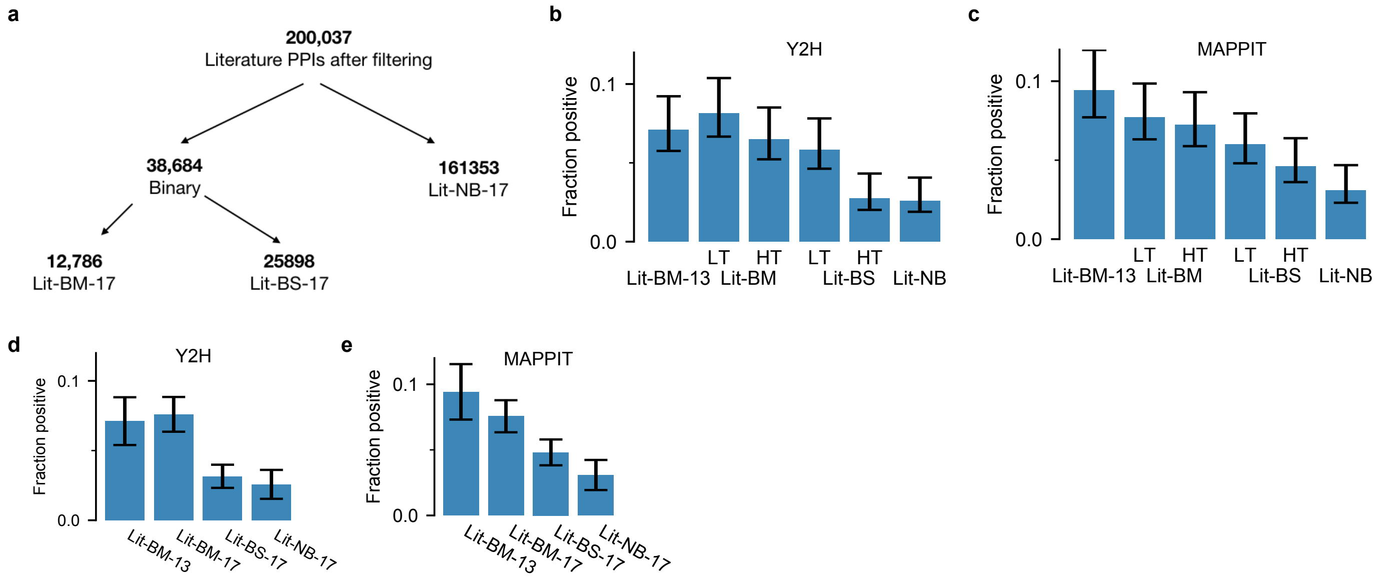


**Extended Data Fig. 9 | Definition of literature-curated PPI datasets.** **a**, Categorization of literature-curated PPIs into distinct subsets based on the experimental methods in which they were detected and the number of pieces of experimental evidence. **b-e**, Results of testing the different categories of literature-curated pairs in Y2H (**b**, **d**) and MAPPIT (**c**, **e**) where the pairs have been further divided into HT - high throughput and LT - low throughput subsets (**b**, **c**). BM: binary multiple; BS: binary singleton; NB: non-binary.

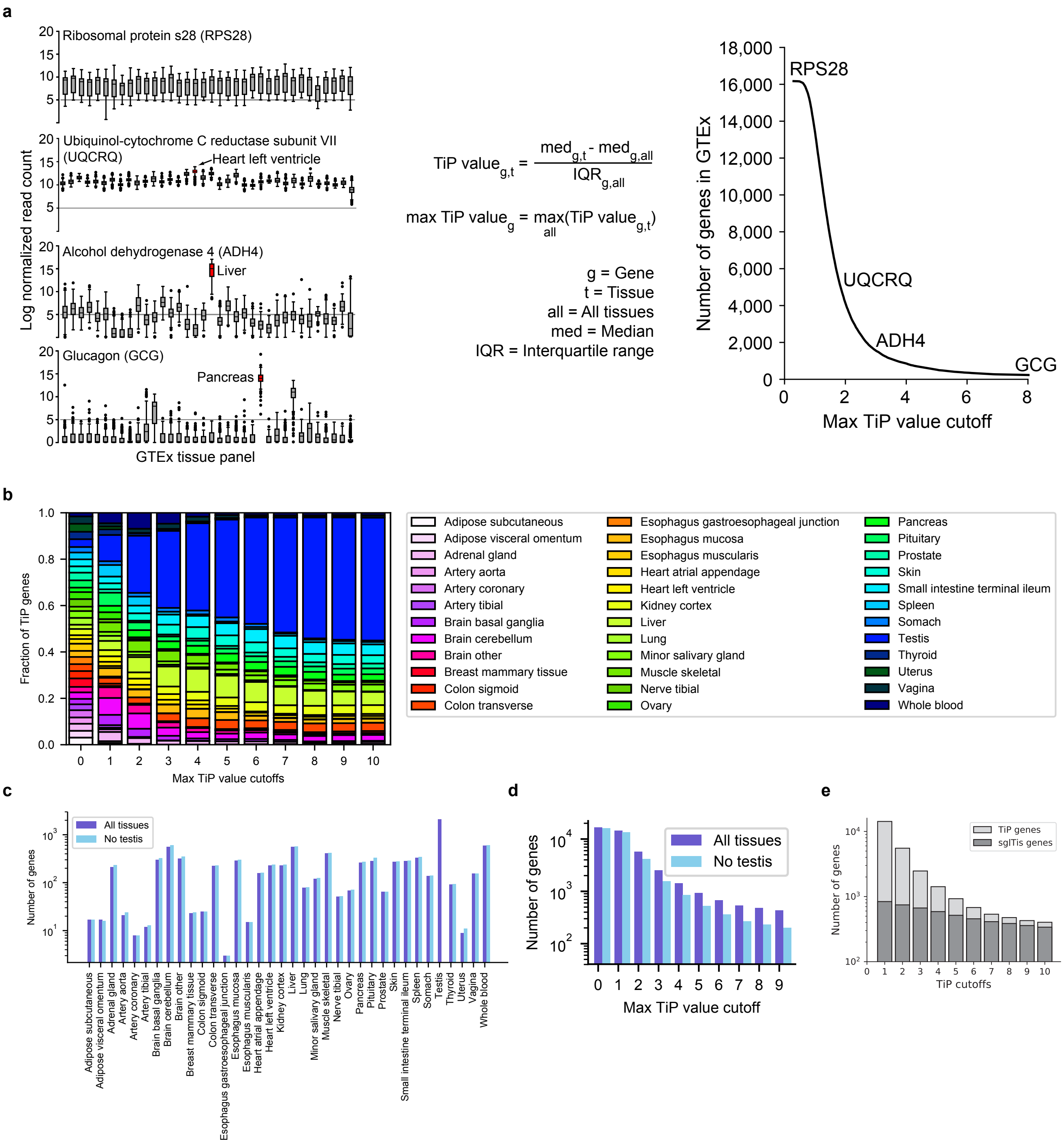


**Extended Data Fig. 10 | Investigation of tissue-preferential expression data.** **a,** Examples of genes displaying different levels of tissue-preferential (TiP) expression across the GTEx tissue panel (left), with boxplots showing median, interquartile range (IQR), and 1.5 x IQR (with outliers). Equation to calculate tissue-preferential expression for every gene-tissue pair and the maximum TiP value for every gene (middle). Number of genes showing tissue-preferential expression for increasing tissue-preferential expression cutoffs (right). **b**, Relative number of TiP genes for every tissue for increasing tissue-preferential expression cutoffs. **c-d**, Differences in number of TiP genes upon removal of testis prior to TiP value calculation per tissue (TiP value cutoff = 2) (**c**) and in total for increasing tissue-preferential expression cutoffs (**d**). **e**, Number of TiP genes and number of TiP genes that are also exclusively expressed in one tissue (sglTis: single tissue) for increasing tissue-preferential expression cutoffs.
